# FtsW exhibits distinct processive motions driven by either septal cell wall synthesis or FtsZ treadmilling in *E. coli*

**DOI:** 10.1101/850073

**Authors:** Xinxing Yang, Ryan McQuillen, Zhixin Lyu, Polly Phillips-Mason, Ana De La Cruz, Joshua W. McCausland, Hai Liang, Kristem E. DeMeester, Cintia C. Santiago, Catherine L. Grimes, Piet de Boer, Jie Xiao

## Abstract

During bacterial cell division, synthesis of new septal peptidoglycan (sPG) is crucial for successful cytokinesis and cell pole morphogenesis. FtsW, a SEDS (Shape, Elongation, Division and Sporulation) family protein and an indispensable component of the cell division machinery in all walled bacterial species, was recently identified in vitro as a new monofunctional peptidoglycan glycosyltransferase (PGTase). FtsW and its cognate monofunctional transpeptidase (TPase) class B penicillin binding protein (PBP3 or FtsI in *E. coli*) may constitute the essential, bifunctional sPG synthase specific for new sPG synthesis. Despite its importance, the septal PGTase activity of FtsW has not been documented *in vivo*. How its activity is spatiotemporally regulated *in vivo* has also remained unknown. Here we investigated the septal PGTase activity and dynamics of FtsW in *E. coli* cells using a combination of single-molecule imaging and genetic manipulations. We show that FtsW exhibits robust activity to incorporate an N-acetylmuramic acid analog at septa in the absence of other known PGTases, confirming FtsW as the essential septum-specific PGTase *in vivo*. Notably, we identified two populations of processive moving FtsW molecules at septa. A fast-moving population is driven by the treadmilling dynamics of FtsZ and independent of sPG synthesis. A slow-moving population is driven by active sPG synthesis and independent of FtsZ’s treadmilling dynamics. We further identified that FtsN, a potential sPG synthesis activator, plays an important role in promoting the slow-moving, sPG synthesis-dependent population. Our results support a two-track model, in which inactive sPG synthase molecules follow the fast treadmilling “Z-track” to be distributed along the septum; FtsN promotes their release from the “Z-track” to become active in sPG synthesis on the slow “sPG-track”. This model explains how the spatial information is integrated into the regulation of sPG synthesis activity and suggests a new mechanistic framework for the spatiotemporal coordination of bacterial cell wall constriction.

## Results

To investigate the role of FtsW in sPG synthesis *in vivo*, we employed a cysteine-modification inactivation assay(1, 2). Based on the homology structure of RodA(3), we generated twelve *ftsW^C^* alleles that each encodes a unique cysteine residue on the periplasmic side of FtsW (Extended Data Fig. 1 and Supplementary Table 1 and 2). Amongst these, we identified FtsW^*I*302*C*^ as a promising candidate for in vivo inactivation with the cysteine-reactive reagent MTSES (2-sulfonatoethyl methanethiosulfonate)(4). Cells expressing FtsW^*I*302*C*^ from the native chromosomal *ftsW* locus grew with a wild-type (WT)-like doubling time and cell morphology in the absence of MTSES but grew into long chains and stopped dividing when treated with MTSES (Extended Data Fig. 2c, Supplementary Movie 1 and 2). WT parental BW25113 cells treated with MTSES did not show any appreciable cell division defect (Extended Data Fig. 2d), indicating that MTSES specifically inhibited the essential function of FtsW^*I*302*C*^. The homology structure of FtsW indicates that I302 resides in periplasmic loop 4 between transmembrane helices 7 and 8, likely near critical residues of the PGTase activity of SEDS proteins(3, 5).

To probe the contribution of FtsW to sPG synthesis in vivo, we labeled new cell wall peptidoglycan (PG) synthesis using an alkyne-modified N-acetylmuramic acid carbohydrate derivative (alkyne-NAM). Unlike fluorescent D-amino acid (FDAA) labels, which are incorporated into the peptide stem of *E. coli* PG through periplasmic exchange reactions(6), alkyne-NAM enters the endogenous cytoplasmic PG biosynthetic pathway and incorporates into newly linked glycan chains(7) composed of alternating units of NAM andN-acetylglucosamine (NAG). Subsequent labeling of the alkyne using a fluorophore-conjugated azide by copper catalyzed azide-alkyne cycloaddition (CuAAC) also known as “CLICK” chemistry allows for visualization and quantification of newly polymerized glycan strands(8).

In WT cells treated with or without MTSES, we observed robust alkyne-NAM labeling (2mg/ml, 30min, Fig. 1a, Methods) at septa. In *ftsW*^*I*302*C*^ cells treated with MTSES (1mM, 30min), we observed a significant reduction in the percentage of cells showing septal labeling above the background level (from 34±4% to 22±4%, *μ*±S.E.M., N>300 cells, three independent repeats, Fig. 1b, orange bars, Supplementary Table 3). Furthermore, in this septal-labeled population of MTSES-treated *ftsW*^*I*302*C*^ cells, the median fluorescence intensity dropped to 70±4% compared to that in untreated cells (Extended Data Fig. 3, Supplementary Table 3). Thus, inhibition of FtsW^*I*302*C*^ activity by MTSES caused a total reduction in septal NAM labeling of 55% (100%–22%/34%×70%, Fig. 1c, orange bar), consistent with a major role of FtsW in septal glycan chain polymerization. The fact that significant NAM incorporation still occurred at septa of MTSES-treated *ftsW*^*I*302*C*^ cells is consistent with previous reports showing that FtsW is not an essential lipid II flippase acting upstream of sPG synthesis(9) and that other PGTases contribute to septal morphogenesis as well(10). The inability of MTSES-treated *ftsW*^*I*302*C*^ cells to complete cell division, even when other PGTases are still active, highlights the essential role of FtsW in successful cell wall constriction.

**Fig. 1.**
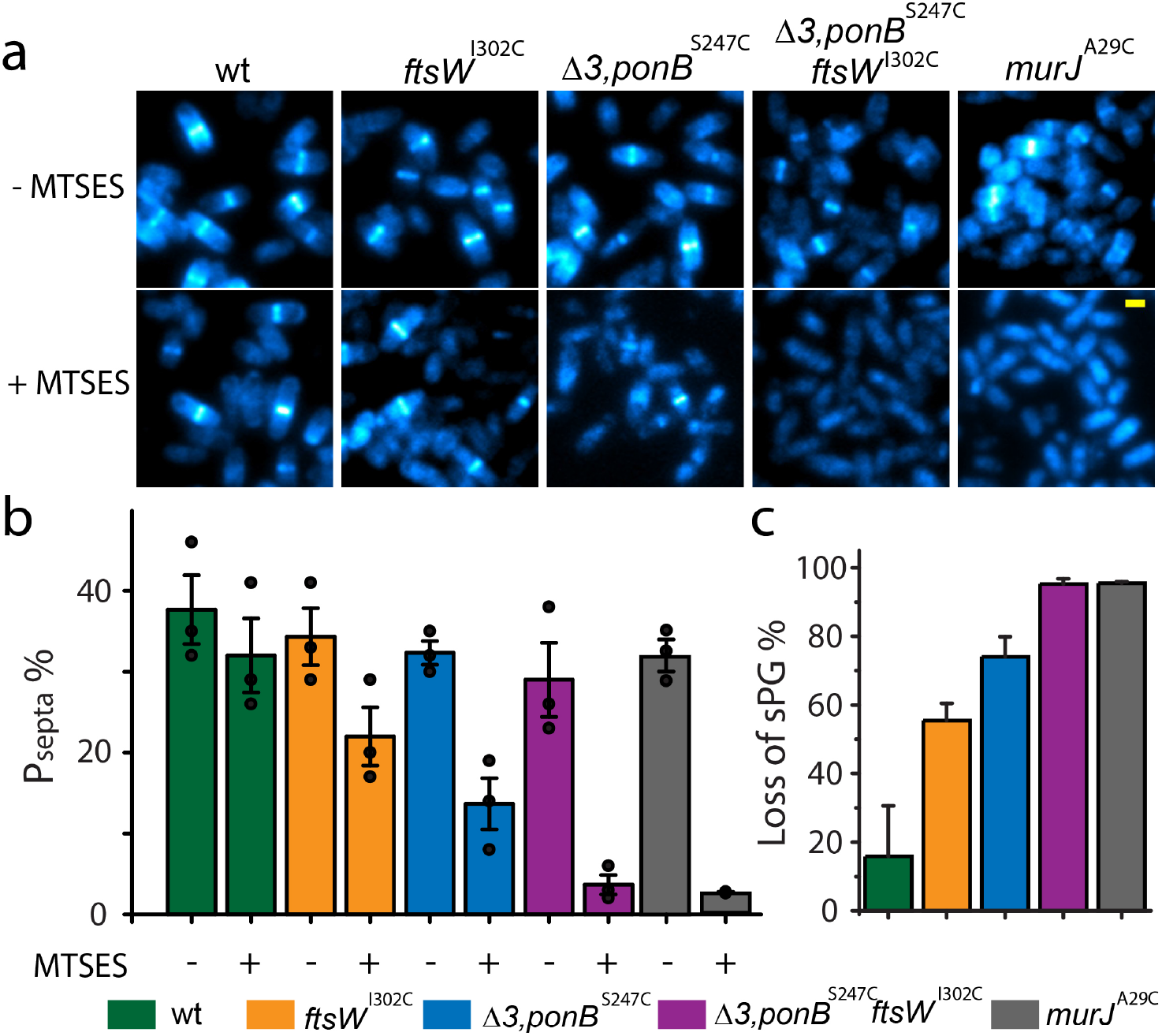
FtsW is an essential septum-specific monofunctional PGTase. **a** Representative images of *E. coli* cells of different strain backgrounds labeled with AF647-conjugated NAM in the absence or presence of MTSES. The contrast of each image is adjusted to allow optimized visualization of septal labeling especially for the Δ3, *ponB^S247C^, ftsW*^*I*302*C*^ and *murJ^A29C^* + MTSES conditions. The absolute intensity is summarized in Supplementary Table 3. Scale bar: 1 *μ*m. **b.** Mean percentage of cells with septal NAM labeling above background level in the absence or presence of MTSES. **c.** Mean percentage of total loss in septal NAM intensity of the five strains due to MTSES. Error bars: S.E.M. of three experimental repeats (dots). Strains used were BW25113 (*wt*), JXY559 (*ftsW*^*I*302*C*^), HC532 (Δ3, *ponB*^*S*247*C*^), JXY564 (Δ3, *ponB*^*S*247*C*^, *ftsW*^*I*302*C*^), and JXY589 (*murJ^A29C^*), all carrying plasmid pBBR1-KU.

To further investigate the relative contributions to sPG polymerization by FtsW and other relevant PGTases in *E. coli*, we introduced the chromosomal ftsW^*I*302*C*^ allele in a Δ3, *ponB*^*S*247*C*^ strain background1 to create a Δ3, *ponB*^*S*247*C*^, *ftsW*^*I*302*C*^ strain. The Δ3, *ponB*^*S*247*C*^ strain lacks the genes for PBP1A, PBP1C and MtgA, and expresses a variant (S247C) of PBP1B (PBP1B^*S*247*C*^, encoded by *ponB*^*S*247*C*^) that, like FtsW^*I*302*C*^, can be inactivated by exposure to MTSES. Untreated Δ3, *ponB*^*S*247*C*^ cells exhibited a similar percentage of labeled septa as WT cells (32±1%, *μ*±S.E.M., N>300 cells, three independent repeats, Fig. 1b, blue bar, Supplementary Table 3). The integrated intensity at septa decreased ~60% (Extended Data Fig. 3, Supplementary Table 3), however, suggesting that the PBP1B^*S*247*C*^ variant may be intrinsically less active or that PBP1A, PBP1C and/or MtgA may also contribute to sPG synthesis under our experimental condition.

When the PGTase activity of PBP1B^*S*247*C*^ in Δ3*ponB*^*S*247*C*^ cells was inhibited by MTSES, the percentage of labeled septa dropped to 14±3% with the median intensity reduced to 65±12% of the untreated level (Fig. 1b, blue bar, Supplementary Table 3), corresponding to a total loss of septal labeling of ~72% (100%–14%/32%×65%, Fig. 1c, blue bar). Simultaneous inactivation of PBP1B and FtsW by MTSES in Δ3, *ponB*^*S*247*C*^, *ftsW*^*I*302*C*^ cells, however, led to a background level of septal NAM labeling indistinguishable from that when the essential Lipid II flippase MurJ(9) was inhibited (Fig. 1b and c, compare purple and gray bars, Supplementary Table 3). Combined with the recent evidence that purified FtsW possesses PGTase activity(11), our results in live *E.coli* cells strongly support the notion that FtsW is an essential septal PGTase *in vivo*.

Previously, we and others showed that FtsZ’s treadmilling dynamics drive the processive movement of FtsW’s cognate TPase (FtsI in *E. coli* and PBP2B in *B. subtilis*) at the septum(12, 13). Such FtsZ-dependent dynamics were proposed to direct the spatial distribution of sPG synthesis complexes and play an important role in septum morphogenesis(12,13). Because a large body of biochemical and genetic studies indicates that FtsW associates with FtsI to form a bifunctional sPG synthase complex(11,14–16), we investigated whether FtsW exhibited similar processive movement at septa using single molecule tracking (SMT).

We constructed a C-terminal fusion of FtsW with the red fluorescent protein TagRFP-T (Supplementary Table 1) and refer to the fusion protein as FtsW-RFP for simplicity. We verified that upon replacement of chromosomal *ftsW* with the *ftsW-rfp* allele (strain JXY422), FtsW-RFP localizes correctly to midcell and supports normal cell division under our experimental conditions (Extended data Fig. 2a, b, and f). To enable single-molecule detection, we expressed FtsW-RFP ectopically at a low level (plasmid pXY349) in the presence of WT FtsW in BW25113 cells. We tracked single FtsW-RFP molecules (Extended Data Fig. 4) at midcell with a 500ms exposure time using wide-field fluorescence microscopy. This slow frame rate allowed us to focus on septum-localized FtsW-RFP molecules by effectively filtering out randomly diffusing molecules along the cylindrical part of the cell body. Using a custom-developed unwrapping algorithm, we decomposed two-dimensional (2D) trajectories of individual FtsW-RFP molecules obtained from the curved cell surfaces at midcell to one-dimensional (1D) trajectories along the circumference and long axis of the cell respectively (Extended Data Fig. 5, Methods).

We found that many single FtsW-RFP molecules displayed directional motions at midcell similar to FtsI(12) (Fig. 2a, Supplementary Movie 3). FtsW-RFP molecules also dynamically transitioned between segments of different speeds and directions (Fig. 2b, Extended Data Fig. 6, Supplementary Movie 4-8). These directional motions were not due to sample drifting as such behaviors were almost absent in fixed cells under the same imaging condition (2.3±0.4%, *μ±* S.E.M., Supplementary Table 6). Some FtsW-RFP molecules displayed confined diffusion and remained largely stationery at the septum (Extended Data Fig. 7a and b, Supplementary Movie S9). Some other FtsW-RFP molecules occasionally entered or left the septum (Movie S6), indicating dynamic exchange of the enzyme between the septum and cell body. Hereafter, we only focus on the dynamics of molecules in the septum where septal cell wall constriction takes place. To quantitatively identify different types of motion of FtsW molecules, we split each trajectory into multiple segments and classified each as stationary or directionally-moving segments based on the total displacement and its associated noise level through statistic means (Fig. 2b, Extended Data Fig. 5e, Methods). Directionally-moving segments were fit to a straight line to extract the directional moving speed *v* (Fig. 2a and b right panels, Extended Data Fig. 5e and 6).

**Fig. 2.**
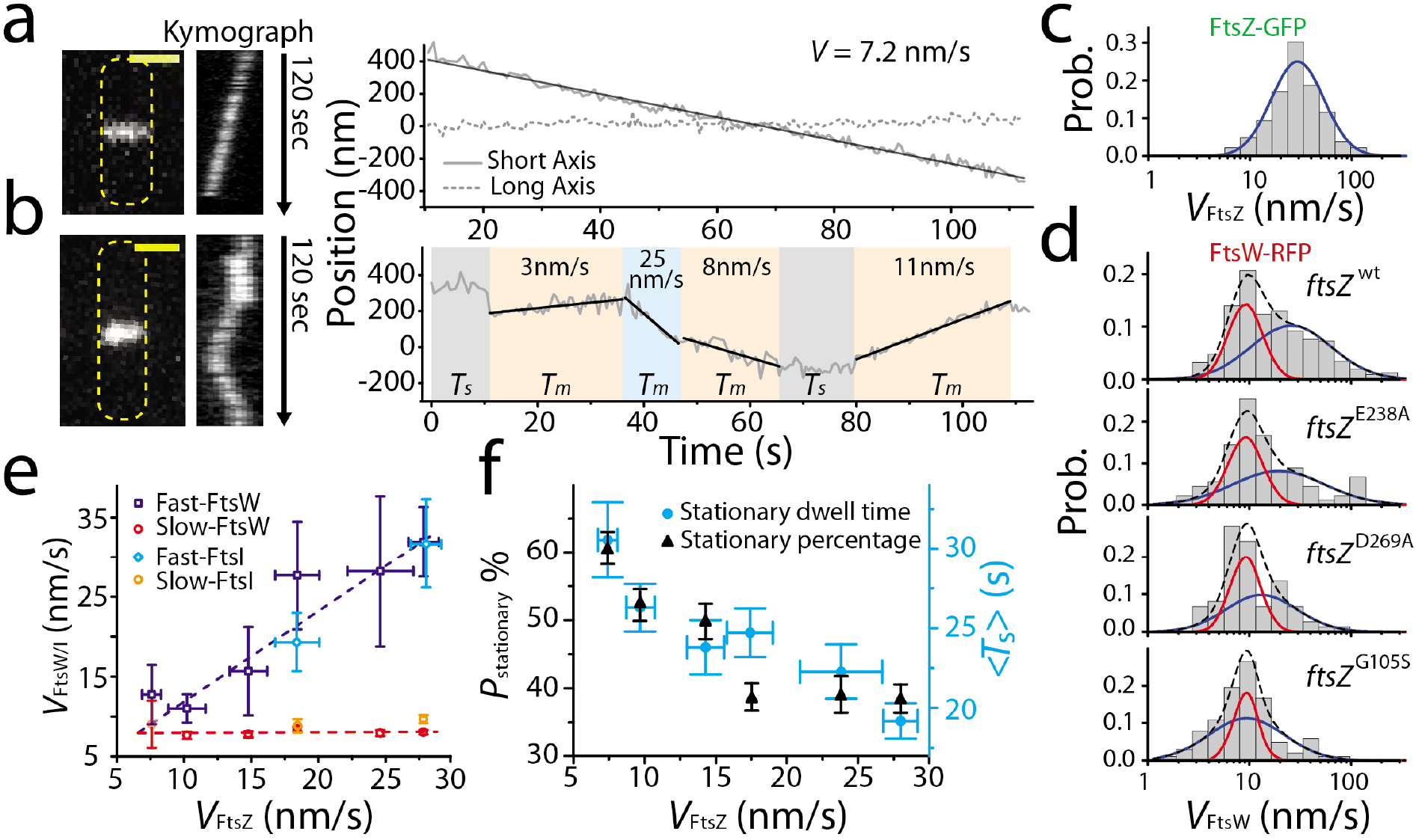
FtsW exhibits two processive moving populations that are differentially dependent on FtsZ’s treadmilling dynamics. **a, b**: Representative maximum fluorescence intensity projection images (left), kymographs of fluorescence line scans at the septa (middle), and unwrapped one-dimensional (1d) positions of the corresponding FtsW-RFP molecule along the circumference (right, solid gray) and long axis (dot gray) of the cell. (a), one processive moving FtsW-RFP molecule and (b), one moving FtsW-RFP molecule that transitioned between different directions and speeds. The trajectory is segmented into different states of slow-moving (beige), fast-moving (pale blue), and stationary (gray). *T_m_*: dwell time of moving segments, *T_s_*: dwell time of stationary segments. Scale bars: 1 *μ*m. **c**. Treadmilling speed distribution of WT FtsZ adapted from (12) overlaid with the single population fit curve (blue). **d**. Speed distributions of directional FtsW-RFP molecules in *wt* and *ftsZ* GTPase mutant strains overlaid with the double population fit curves (slow-moving population in red, fast-moving population in blue, and the overall fit curve in black dash lines). Note that the red population remained largely unchanged while the blue population gradually moved to slower speeds in FtsZ GTPase mutants. **e**. Decomposed mean speed of the fast population (blue: FtsW, cyan: FtsI) is correlated with FtsZ’s treadmilling speed while the speed of the slow-moving population (red: FtsW, orange: FtsI) is independent on FtsZ’s treadmilling speed. **f**. Percentage (dark triangles) and mean dwell time (cyan dots) of stationary FtsW molecules increased with reduced FtsZ treadmilling speed (left to right).

Notably, the speeds of all directionally-moving FtsW-RFP molecules showed a much wider distribution (Fig. 2d, top panel, N=315 segments) than the distribution of FtsZ’s treadmilling speed(12) (Fig. 2c). The FtsW-RFP’s speed distribution was best fit by the sum of two populations, one fast and one slow (Fig. 2d, blue and red curves, Extended Data Fig. 8, Supplementary Table 4 and 9, Methods), distinct from a single fast population as that was seen for FtsZ’s treadmilling speed distribution (Fig. 2c, blue curve). The fast-moving population (62.6±11.2%, *μ*±S.E.M.) of FtsW-RFP displayed a mean speed of 31.9±4.4 nm/s, (*μ*±S.E.M., Fig. 2d top panel, blue curve and 2e, Supplementary Table 4), close to the average FtsZ treadmilling speed we previously measured(12) (28.0±1.2 nm/s, *μ*±S.E.M.). The slow-moving population (37.4±11.2%, *μ*±S.E.M.) of FtsW-RFP molecules had an average speed of ~8nm/s (8.0±0.3nm/s, Fig. 2d, top panel, red curve, Supplementary Table 4). The existence of, and transition between, the two different types of directional movements could also be directly observed in many individual FtsW-RFP trajectories (Fig. 2b, Extended Data Fig. 6). The essential sPG transpeptidase FtsI, which likely forms a cognate complex with FtsW, exhibits a statistically indistinguishable speed distribution compared to that of FtsW (p=0.32, K-S test), with similar two directional-moving populations (Extended Data Fig. 9, 46.1±19.9% and 31.2±5.6 nm/s for the fast moving population, 53.9±19.9% and 9.8±1.1 nm/s for the slow-moving population, *μ*±S.E.M., n=92 segments, Supplementary Table 5).

To investigate how the two directionally moving populations of FtsW respond to FtsZ’s treadmilling dynamics, we performed SMT of FtsW-RFP in five strains with mutations that progressively lower FtsZ’s GTPase activity and treadmilling speed (*ftsZ*^*E*238*A*^, *ftsZ*^*E*250*A*^, *ftsZ*^*D*269*A*^, *ftsZ*^*G*105*S*^ and *ftsZ*^*D*158*A*^, Fig. 2d, Supplementary Table 4). As we previously also observed for FtsI(12), the average speed of all moving FtsW-RFP molecules decreased with the tread-milling speed of FtsZ in these mutant strains (Supplementary Table 4). However, decomposing the speed distribution of FtsW-RFP into two moving populations in each FtsZ GTPase mutant (Fig. 2e, Methods, Extended Data Fig. 8) revealed that only the speed of the fast-moving population is highly correlated with FtsZ’s treadmilling speed (Fig. 2e, blue), whereas the slow-moving population maintained a relatively constant speed around 8nm/s independent of FtsZ’s treadmilling speed (Fig. 2e, orange, Supplementary Table 4). A similar differential response of FtsI was observed in the *ftsZ*^*E*250*A*^ background under the same experimental condition (19.2±3.4nm/s and 9.1±1.8nm/s for the fast and slow-moving populations respectively, *μ*±S.E.M, n=45 segments, Fig. 2e, cyan and orange markers, Extended Data Fig. 9, Supplementary Table 5). This observation supports the idea that FtsW and FtsI likely move together as part of one sPG synthase complex. The differential responses of the two moving populations of FtsW and FtsI to FtsZ’s treadmilling dynamics suggest that the fast-moving population is driven by FtsZ treadmilling dynamics as we showed previously(12), while the slow-moving population is not. Note that a recent study showed that in *S. pneumoniae* the sPG synthesis complex, FtsW-PBP2x, only exhibited one single, slow-moving population that is FtsZ-independent(17).

In addition to the two directional moving populations, we investigated how the stationary population of FtsW-RFP responded in FtsZ GTPase mutants. Using the mean squared displacement (MSD) analysis, we found that stationary FtsW-RFP molecules in WT FtsZ cells were confined in a region of ~100nm in length (95.5±6.1nm, *μ*±S.E.M., N=179 segments) with a very small diffusion coefficient (D=0.0007±0.0002*μ*m^2^/s, *μ*±S.E.M., Extended Data Fig. 7c). We confirmed that this level of confinement was not due to our experimental uncertainty in SMT, as fixed cell SMT showed a much smaller mean square displacement (Extended Data Fig. 7c). Because the confinement length is on par with the average length of FtsZ filaments in the FtsZ-ring (80-160nm) (18, 19), it is possible that some of these stationary FtsW molecules are bound to the middle of FtsZ filaments. Consistent with this possibility, we observed that the average dwell time and percentage of the stationary FtsW-RFP population increased as the treadmilling speed of FtsZ decreased in GTPase mutant strains (Fig. 2f, Supplementary Table 4), in line with the observed increase in the dwell time of FtsZ monomers and FtsZ filament length in FtsZ GTPase mutants in a recent study (20). Interestingly, the average dwell time of stationary FtsW-RFP in wt cells (19.2±1.1s, Fig. 2f, Supplementary Table 4) is significantly longer than the lifetime of FtsZ monomers (8.1±0.5s) (20), indicating that other mechanism(s) may also be at play to maintain this stationary population of FtsW-RFP, as we will further describe below.

Given our results so far that the overall sPG synthesis activity is not affected in FtsZ GTPase mutants in *E. coli* (12, 21), it is likely that the fast-moving population of FtsW and FtsI is inactive in sPG synthesis, but dependent on FtsZ’s treadmilling to be transported to different septal positions. We recently have shown that these sPG enzyme molecules could follow FtsZ’s fast treadmilling dynamics by continuously tracking the shrinking end of FtsZ filaments through a Brownian Rachet mechanism (22). The FtsZ-independent, slow-moving population, however, may be active in sPG synthesis since their speed is independent of FtsZ. To examine this possibility, we investigated the speed distributions of FtsW-RFP molecules under conditions of altered sPG synthesis activity.

We first examined the movement of FtsW-RFP molecules when its PGTase activity is enhanced or inhibited (Fig. 3a-c). To examine FtsW’s motion under enhanced sPG synthesis activity conditions, we tested a superfission (SF) allele of *ftsW* (*ftsW*^*E*289*G*^) that confers a short-cell phenotype and alleviates the need for FtsN, an otherwise essential positive regulator of sPG synthesis (Extended Data Fig. 10). SMT of the SF variant (FtsW^*E*289*G*^-RFP) in *ftsW*^*E*289*G*^ cells again revealed two populations, one fast and one slow as its WT parental strain TB28 (Fig. 3a, blue and red curves), but the fast-moving population was reduced from 36.4±7.6% to 20.9±5.6% (Fig. 3f, Supplementary Table 6). The mean speed of each population did not change significantly compared to that of the WT strain (Supplementary Table 6). Accordingly, the average speed of all directionally moving FtsW^*E*289*G*^-RFP molecules in *ftsW*^*E*289*G*^ cells was significantly slower than that of FtsW-RFP in WT parental TB28 cells (13.9±1.1 vs 18.8±1.6nm/s, Fig. 3f, blue, Supplemental Table 6). These results are consistent with the hypothesis that the slow-moving population of FtsW is coupled to sPG synthesis activity and that the SF FtsW^*E*289*G*^ variant increased this slow-moving population significantly.

**Fig. 3.**
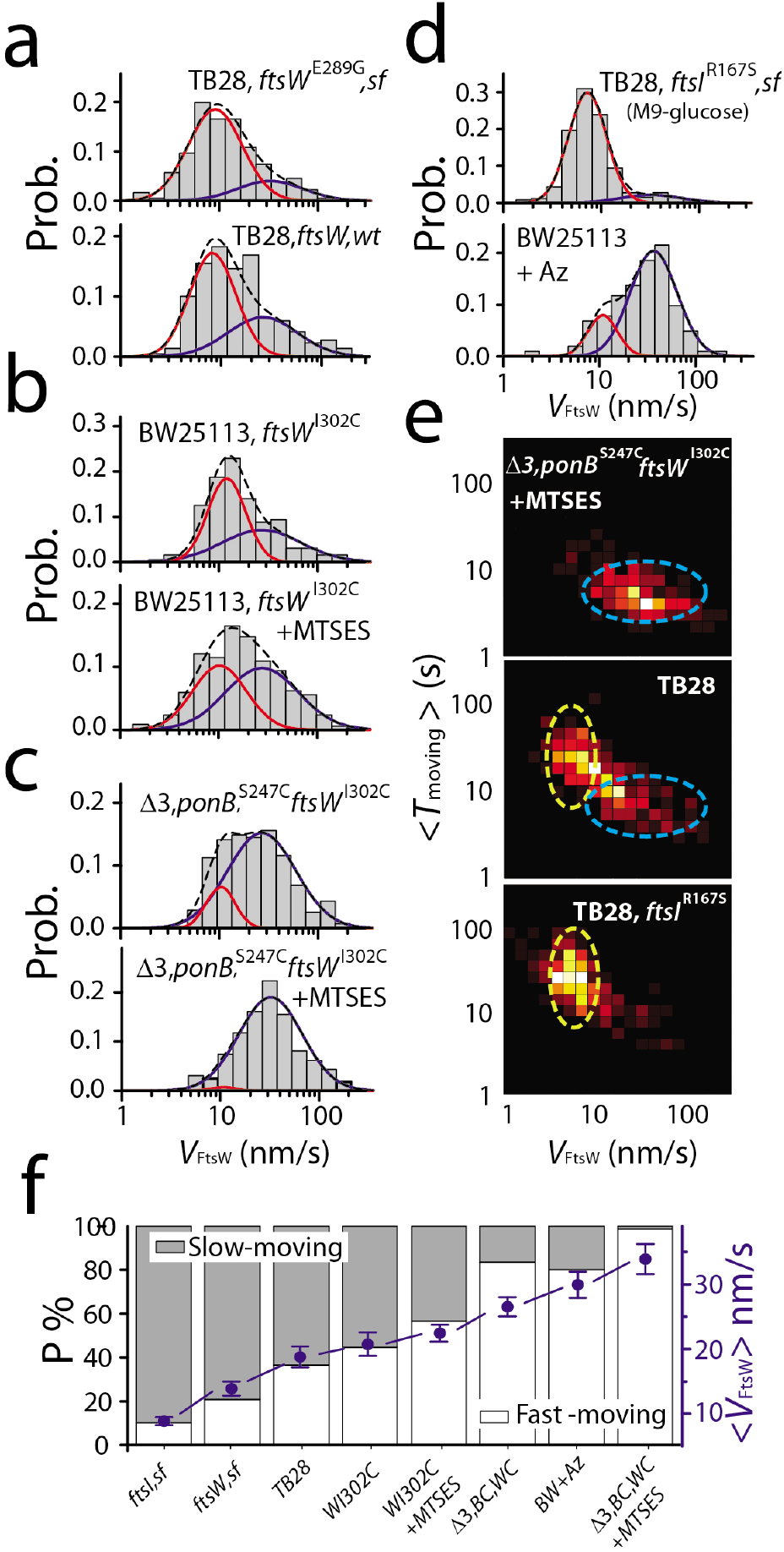
The slow-moving population of FtsW increases with enhanced sPG synthesis activity and depletes with reduced sPG synthesis activity. **a-d.** Histograms of directional moving speeds of FtsW-RFP (gray bar) overlaid with two population fitting curves (slow-moving population in red, fast-moving population in blue, and overall fit curve in black dash lines). **(a)** The superfission variant FtsW^*E*289*G*^-RFP in the TB28, *ftsW*^*E*289*G*^ strain background (top) shows increased slow-moving population (red curve) compared to that of wildtype FtsW-RFP in the parental TB28 strain background. **(b)**. FtsW^*E*289*G*^-RFP in JXY559 (*ftsW*^*E*289*G*^) in the absence (top) of MTSES shows increased slow-moving population (red curve) compared to that in the presence of MTSES where FtsW activity is inhibited (bottom). **(c)**. FtsW^*E*289*G*^-RFP in JXY564 (Δ3, *ponB*^*S*247*C*^, *ftsW*^*I*302*C*^) in the presence of MTSES exhibits nearly completely abolished slow-moving population (bottom) compared to that in the already significantly diminished slow-moving population in the absence of MTSES (top). **(d)** Wildtype FtsW-RFP in the FtsI su-perfission strain TB28, *ftsI*^*R*167*S*^ background in M9-glucose medium (top) exhibits the most enhanced slow-moving population compared to that in the BW25113 (*wt*) strain treated with FtsI-inhibitor Aztreonam. **e.** 2D-heatmap of the moving speed and dwell time of directional moving FtsW (or FtsW^*I*302*C*^)–RFP molecules in normal growth condition (M9-glucose, middle), all PGTase inhibited condition (top), and FtsI superfission strain (bottom). The slow-moving population with long dwell time (yellow oval) and the fast-moving population with short dwell time (blue oval) are marked with circles as a guide for the eyes. **f.** Percentage of the slow- (gray bar) and fast- (white bar) moving population under all the conditions in a-d. Average moving speeds under all the conditions are plotted in blue dots.

Next, we inhibited partially or completely septal glycan strand polymerization using MTSES on two strains we describe in Fig. 1. We first tracked the movement of FtsW^*I*302*C*^-RFP molecules in the BW25113 *ftsW*^*I*302*C*^ background (JXY559/pAD004) in the absence and presence of MTSES. In the absence of MTSES, we again observed two directionally moving populations of FtsW^*I*302*C*^-RFP, one fast at 33.8±2.7nm/s (44.6±7.5%) and one slow at 10.7±0.9nm/s (55.4±7.5%). The average speed is at 20.7±1.8nm/s (*μ*±s.e.m., N=192 segments, Fig. 3b, top panel, Supplementary Table 6). In the presence of MTSES, the slow-moving population of FtsW^*I*302*C*^-RFP was reduced to 43.4±9.1% (Fig. 3b, bottom panel, *μ*±s.e.m., N=339 segments, Supplementary Table 6). The average speed increased to 22.5±1.3 nm/s (*μ*±s.e.m., Fig. 3f, Supplementary Table 6).

Since MTSES specifically blocks FtsW^*I*302*C*^-dependent sPG synthesis, we reasoned that the remaining slow-moving population of FtsW could be driven by sPG synthesis from PBP1A, 1B, 1C, and MtgA (Fig. 1). To examine this possibility, we tracked FtsW^*I*302*C*^-RFP molecules in the Δ3, *ponB*^*S*247*C*^, *ftsW*^*I*302*C*^ background (JXY564/pAD004). In this background, even in the absence of MTSES, FtsW^*I*302*C*^ already displayed a significantly reduced slow-moving population (16.5±6.2%, *μ*±s.e.m., N=276 segments, Fig. 3c) accompanied by an increased fast moving population compared to WT TB28 cells (63.6±7.6%, *μ*±s.e.m., N=219 segments, Fig. 3a, bottom panel). These changes are consistent with the reduced alkyne-NAM incorporation in this strain background (Extended Data Figure. 3), suggesting that the decrease of the overall PGTase activity indeed depletes the slow-moving population. The average speed also increased to 26.3±1.5nm/s. Further abolishing all PGTase activity by 0.1mM MTSES (~99% reduction, Supplementary Table 3) caused a near complete depletion of the slow-moving population to 1.2±2.7%, (Fig. 3c bottom panel and f), with the fast-moving population speed remaining at 32.1±1.5nm/s (Supplementary Table 6).

To ensure that the depletion of slow-moving FtsW molecules is not caused by altered FtsZ treadmilling dynamics under these conditions, we measured FtsZ dynamics in those strains in the presence or absence of MTSES.

FtsZ’s treadmilling speed distribution remained unchanged (Extended Data Fig. 11b, Supplementary Table 6), supporting the hypothesis that the slow-moving population of FtsW is related to sPG synthesis activity rather than to FtsZ treadmilling.

To further verify that the slow-moving population of FtsW-RFP is related to active sPG synthesis, we monitored the movement of FtsW-RFP in cells expressing a superfission variant of FtsI (FtsI^*R*167*S*^). FtsI^*R*167*S*^ is similar to FtsW^*E*289*G*^ in alleviating the essentiality of FtsN but does so only partially (Extended Data Fig. 12, Table S7). In *ftsI*^*R*167*S*^ cells, a major population of FtsW-RFP (90.7±2.1%,) moved slowly at 6.4±0.2nm/s and a minor population moved fast at 31.6±2.0 nm/s (9.3±2.1%, N=255 segments, Fig. 3d, top panel, Supplementary Table 6). The average speed is drastically reduced to 8.8±0.6nm/s (Fig. 3f, Supplementary Table 6). When we applied FtsI-specific β-lactam Aztreonam to inhibit its transpeptidase (TPase) activity (1*μ*g/mL, 30 min)(23) in WT BW25113 cells, we observed that the slow-moving population of FtsW-RFP was depleted from 37.4±11.2% to 19.8±7.7% (Fig. 3d, red curve), whereas the fast-moving population increased with a similar speed distribution as FtsZ’s treadmilling dynamics (Fig. 3d, blue curve, Extended Data Fig. 11b). The average speed of FtsW-RFP became 29.9±2.0nm/s (N=102 segments). These results not only support the hypothesis that the slow-moving population of FtsW depends on active sPG synthesis, but also indicate that the crosslinking of sPG by FtsI’s TPase activity is equally important as the polymerization of glycan strands to drive the progressive movement of the slow population of the sPG synthase.

Most interestingly, we found that the two moving populations of FtsW are also separated in the time domain. We measured the dwell time that individual FtsW-RFP molecules spent at a constant speed (as depicted in Fig. 2b) and plotted the corresponding two-dimensional (2D)-histogram of the moving speed and dwell time in Fig. 3e. We found that the slow-moving population (~8nm/s) is associated with a relative long dwell time (~25sec, yellow circle in Fig. 3e middle panel) and the fast-moving population (~30nm/s) with a significantly shorter one (~6sec, blue circle in Fig. 3e). In the sPG synthesis inhibited (Δ3, *ponB*^*S*247*C*^, *ftsW*^*I*302*C*^ +MTSES) condition, the fast-moving, short-lived population (5.8±3.2 sec, *μ*±s.d.) became dominant (Fig. 3e, top panel, Extended Data Fig. 13), while the slow-moving, long-lived population (25.3±15.6 sec, *μ*±s.d.) dominated in the superfission *ftsI*^*R*167*S*^ background (Fig. 3e bottom panel, Extended Data Fig. 13). These observations further confirmed that the two moving populations of FtsW-RFP molecules are distinct from each other.

One factor controlling the sPG synthesis rate is the availability of cell wall precursors. In *E. coli* under balanced growth, the cellular level of precursors limits cell growth and cell wall constriction rates(24, 25). We reasoned that if the slow-moving population of FtsW were indeed driven by sPG synthesis activity, its size and even its speed might be modulated by the level of available PG precursors. To test this possibility, we first depleted precursor Lipid II in WT BW25113 cells by blocking the essential enzyme MurA using Fosfomycin(26). We found that in cells treated with Fosfomycin (50*μ*g/ml, 30min), the slow-moving population of FtsW was significantly diminished to ~8% (7.9±9.3%, N=239 segments, Fig. 4a and b, Supplementary Table 6) while the fast-moving population dominated with unchanged moving speed.

**Fig. 4.**
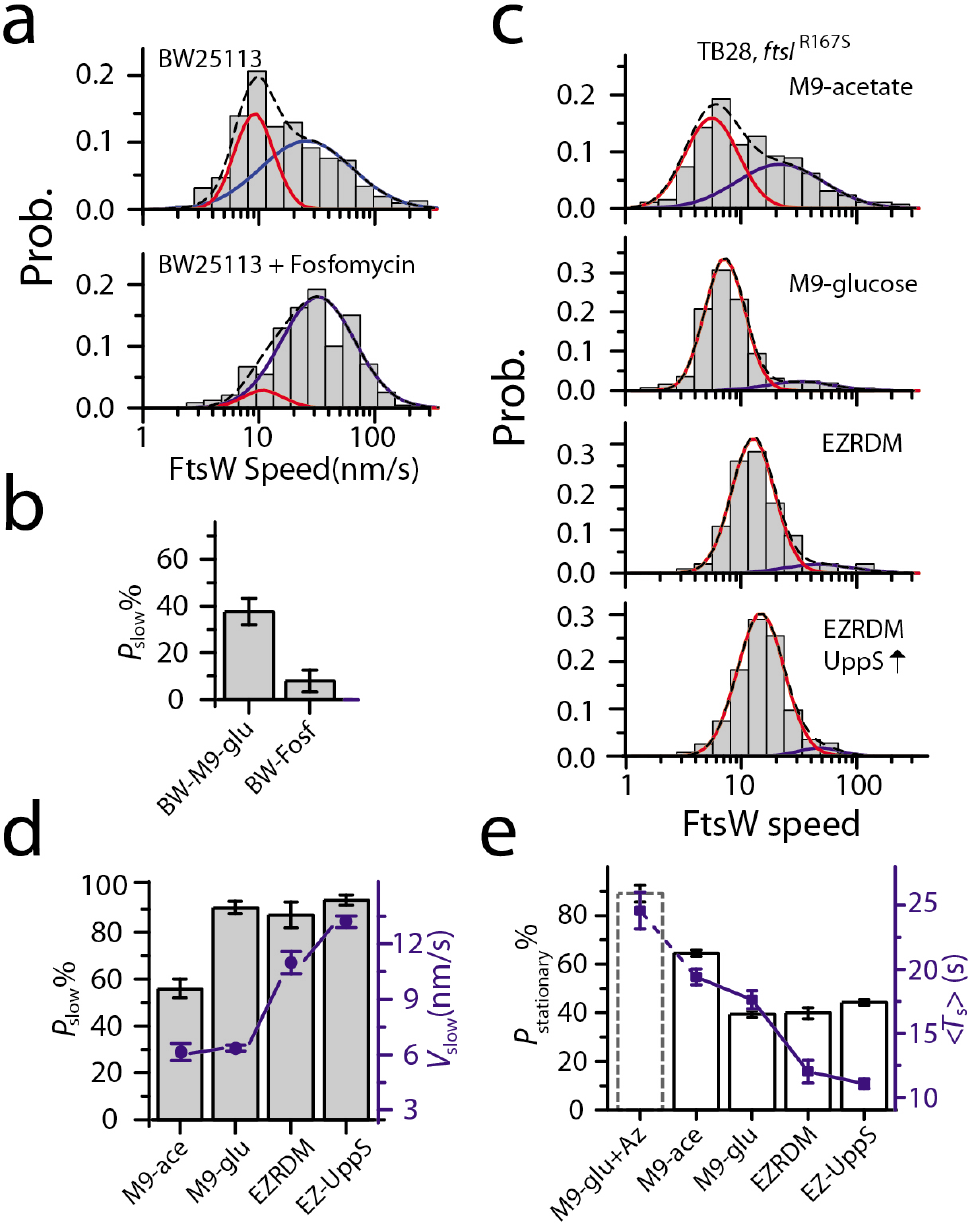
Enhanced growth conditions promote the slow-moving population of FtsW. **a, c.** Histograms of directional moving speeds of FtsW-RFP (gray bar) overlaid with slow-(red) and fast-(blue) moving population fit curves, and the overall fit curve (black dash). **(a)** FtsW-RFP in BW25113 cells in M9-glucose medium (top, replot from Fig. 2d) and treated with Fosfomycin to deplete the lipid II precursor pool (bottom). **(b)** The slow-moving population of FtsW-RFP decreases upon depletion of lipidII (Fosfomycin treatment). **c.** FtsW-RFP in FtsI superfission strain TB28, *ftsI*^*R*167*S*^ grown in M9-acetate, M9-glucose, EZRDM, or in EZRDM medium and with UppS overproduction (top to bottom). **d.** The percentage (gray bar) and average speed (blue dot) of the slow-moving population increases in media with enhanced growth conditions. **e.** Percentage (white bars) of stationary FtsW-RFP molecules decreases and reaches a plateau with enhanced growth conditions whereas and average dwell time (blue squares) continues to decrease. BW25113 *wt* cells treated with Aztreonam treatment (dashed bar, M9-glu+Az) is shown as a comparison. The corresponding constriction rates for all the growth conditions are listed in Supplementary Table 8.

Next, we used M9 minimal growth medium containing acetate or glucose as the carbon source or EZ rich defined medium (EZRDM) to vary the available levels of PG precursors in the superfission *ftsI*^*R*167*S*^ background (strain PM6). We measured the corresponding cell wall constriction rates using an mNeonGreen-ZapA fusion protein to confirm that cells growing in these different media exhibited different sPG synthesis activities. Indeed, the constriction rate increased from 4.9±1.1nm/min (N=5 cells) in M9-acetate, to 17.2±1nm/min (N=15 cells) in M9-glucose, and finally to 37.9±2.0nm/min (N=16 cells) in EZRDM medium (Method, Extended Data Fig. 14, Supplementary Table 8). We then monitored the motion of FtsW-RFP under these different growth conditions. As shown in Fig. 4c, the fast-moving population of FtsW, originally diminished in the M9-glucose medium in the superfission variant background (Fig. 3d, top panel, replotted in the second panel of Fig. 4c for a comparison), re-appeared under the M9-acetate growth condition (44.4±4.8%, N=260 segments, Fig. 4c top panel). This result, together with our observation using Fosfomycin, strongly supports our hypothesis that the percentage of slow-moving FtsW molecules is governed by sPG synthesis activity, here determined by the level of cell wall precursors.

In rich growth medium (EZRDM), the percentage of slow-moving FtsW-RFP molecules in PM6 (*ftsI*^*R*167*S*^) cells stayed around 90% (87.5±5.5%, Fig. 4d), similar to that in M9-glucose. However, the average speed of the slow-moving population of FtsW-RFP accelerated to 11.0±0.6nm/s (*μ*±s.e.m., N=92 segments, Fig. 4c second and third panel, 4d, blue) in EZRDM, ~72% higher than in M9-glucose (6.4±0.2nm/s). This faster speed of FtsW-RFP is still significantly slower than that of FtsZ’s treadmilling speed under the same growth condition (Extended Data Fig. 11c, Supplementary Table 6), consistent with the expectation that this population is not coupled to FtsZ’s treadmilling dynamics. Note that growth of wt BW25113 cells in EZDRM also led to an increased speed of the slow-moving population of FtsW-RFP (11.1±0.2 nm/s, *μ*±s.e.m., n=910 segments, Supplementary Table 6), indicating that this medium effect is not specific to the superfission FtsI variant in PM6 cells.

Finally, to further upregulate Lipid II precursor levels in PM6 (*ftsI*^*R*167*S*^) cells, we grew these cells in EZDRM and induced the expression of the undecaprenyl pyrophosphate synthetase UppS from a plasmid(9). Under this condition, we observed a further increase in the slow-moving speed of FtsW-RFP to 13.2±0.3nm/s (*μ*±s.e.m., N=513 segments, Fig. 4c bottom panel and 4d, blue). Taken together, these results strongly suggest that the percentage of the slow-moving population reflects the relative amount of active FtsW molecules engaged in sPG synthesis, and its speed likely reflects the *in vivo* sPG polymerization rate at the individual enzyme molecule level.

In the above experiments, we also observed changes in the percentages and dwell times of the stationary population of FtsW-RFP (Fig. 4e). The percentage of stationary molecules decreased as the expected cell wall precursor levels increased and reached a plateau at ~40% under the M9-glucose, EZRDM and UppS-overexpression conditions (Fig. 4e, white bars). The average dwell time of stationary FtsW-RFP molecules, however, continued to decrease to ~10s (Fig. 4e, blue squares), approaching the mean lifetime of FtsZ monomers in the Z-ring (8.1 sec)(20).

These results, together with those from the experiments of FtsZ GTPase mutants (Fig. 2f), suggest that at least two subpopulations of stationary FtsW molecules may exist. One likely represents molecules bound to internal subunits of FtsZ filaments, and their dwell times and abundance are determined by the length and treadmilling speed of FtsZ filaments (Fig. 2f). The other subpopulation may represent FtsW molecules that are poised at sPG synthesis sites waiting for lipid II to become available. This possibility is supported by our observations that the percentage and mean dwell time of this subpopulation indeed shrank as the level of cell wall precursors increased in rich medium (M9-glucose and EZRDM) or with upregulated UppS expression (Fig. 4d, Extended Data Fig. 7d and e, Supplementary Table 6).

We further reasoned that Aztreonam treatment could substantially enhance the stationary population. Under this condition FtsW engages in a futile sPG synthesis cycle in which it continues to polymerize glycan strands, but these are blocked from being crosslinked by FtsI and subsequently degraded(27). Therefore, the lack of FtsI’s crosslinking activity may prevent directional movement of the enzyme while the continuous polymerization activity of FtsW prevents its dissociation from glycan strands. Consistent with this expectation, the dwell time and percentage of the stationary population of FtsW-RFP in Aztreonam-treated cells increased to 24.6±1.4 sec and 88.9±3.4% respectively (n=347 segments, Fig. 4e, Supplementary Table 6). Furthermore, the MSD curve of stationary FtsW-RFP molecules became indistinguishable from that of the fixed sample (Extended Data Fig. 7c, green), suggesting they also become confined to a smaller region after Aztreonam treatment. Note that previously a stationary, inactive, and PG-bound state has been reported for *E. coli* PBP2, the counterpart of FtsI in the cell elongation machinery(28).

Our results so far demonstrated that two processive moving populations of FtsW exist *in vivo*. The fast-moving population is driven by FtsZ’s treadmilling dynamics but inactive in sPG synthesis, whereas the slow-moving population is most likely active and driven by sPG synthesis. The presence of active and inactive FtsW populations raises an interesting question: Do division proteins involved in controlling the initiation and progression of cell wall constriction also determine the partitioning of these two populations? Previous studies in *E. coli* have shown that when FtsW and FtsI are first recruited to the division site by a complex of the FtsB, FtsL and FtsQ proteins (FtsQLB), they are kept in an inactive state until FtsN, the last essential division protein to accumulate at the site, activates the sPG synthesis complex (29, 30). Thus, FtsN may play an important role in triggering the transition of FtsW from the fast-moving (FtsZ-dependent) to slow-moving (sPG synthesis-dependent) state.

To test this hypothesis, we used an FtsN-depletion strain (EC1908) (31) wherein the chromosomal *ftsN* gene is controlled by the *araBAD* regulatory region. This strain grows and divides normally in the presence of 0.2% of arabinose (Extended Data Fig. 15). After overnight growth (~15 hours) in M9-glucose medium without arabinose, the average cellular FtsN level was depleted to ~44% of that in WT cells (Extended Data Fig. 15) and many cells grew into long filaments with shallow constrictions (Fig. 5a, bottom panel). We tracked FtsW-RFP molecules at shallow constriction sites in these filamentous cells. As expected, the slow-moving population of FtsW-RFP was reduced significantly from 63.6±7.6% to 41.7±14.7% (Fig. 5b, third and bottom panels, orange curves, Supplementary Table 6), and the increased fast-moving population of FtsW-RFP molecules moved at an average speed of 30.1±3.9nm/s (*μ*±s.e.m., N=398 segments), similar to that of FtsZ treadmilling. This result suggests that FtsN plays a role in promoting the slow-moving, or active, population of FtsW.

**Fig. 5.**
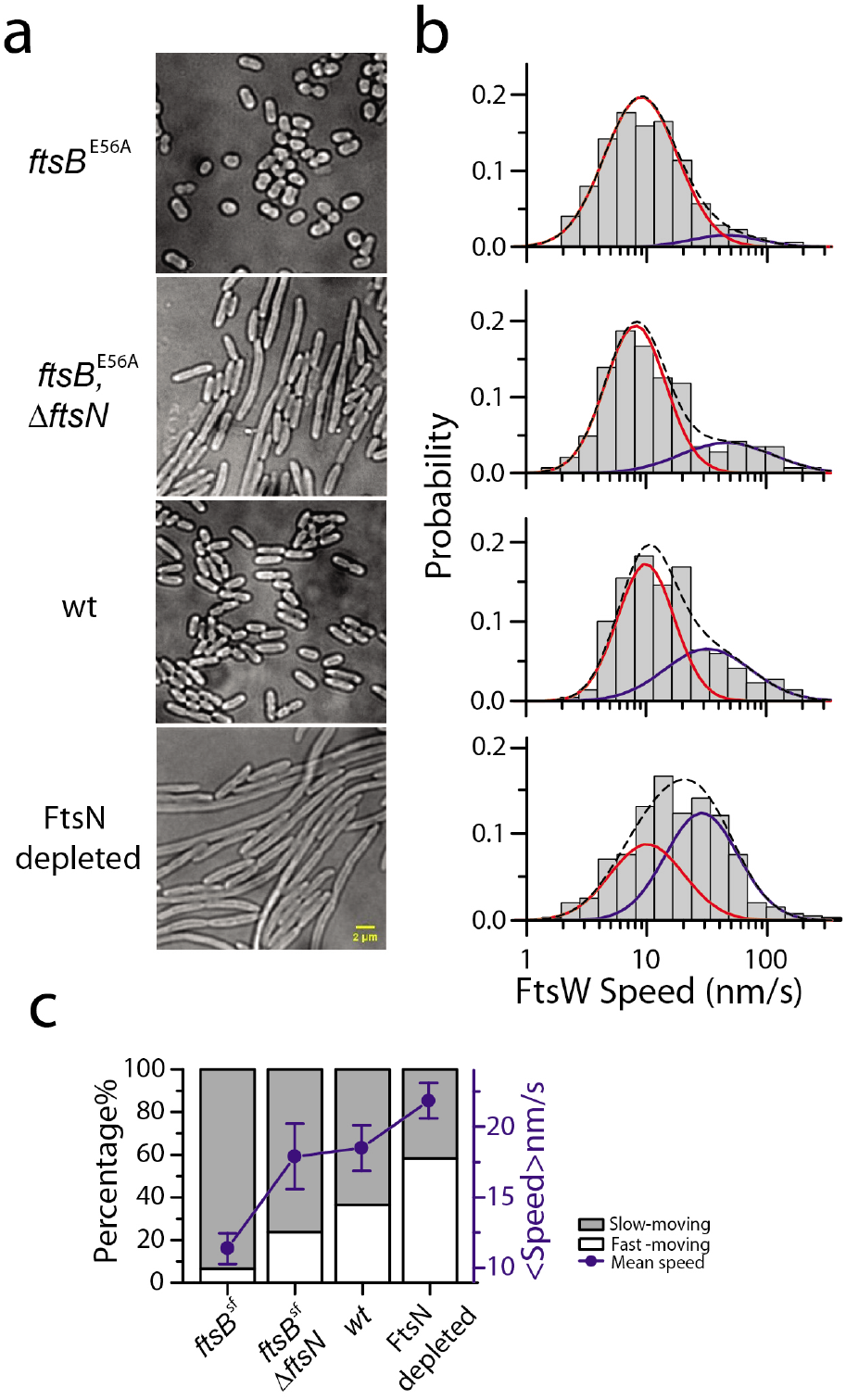
FtsN plays an important role in promoting the slow-moving population of FtsW. **a.** Bright-field images of superfission *ftsB*^*E*56*A*^ (BL167), *ftsB*^*E*56*A*^ Δ *ftsN* (BL173), WT (TB28), and FtsN-depleted cells (EC1908). Scale bar: 2 *μ*m. **b.** Histograms of directional moving speeds of FtsW-RFP (gray bar) overlaid with slow- (red) and fast-moving (blue) population fit curves, and the overall fit curve (black dash) in cells corresponding to a on the left (in M9-glucose medium). **c.** Average FtsW-RFP moving speeds (blue dots) and percentage of the slow-(gray bar) and fast-(white bar) moving populations under the conditions in a-b.

Next, to assess the effects of FtsN on FtsW dynamics under conditions were FtsN is no longer essential, we tracked the movements of FtsW-RFP in a *ftsB*^*E*56*A*^ superfission strain (BL167) that still produces FtsN, and also in strain BL173 (*ftsB*^*E*56*A*^, Δ*ftsN*) that lacks FtsN completely. The *ftsB*^*E*56*A*^ superfission allele causes cells to initiate sPG synthesis earlier in the division cycle than normal, leading to a small-cell phenotype (29) (Fig. 5a, top panel). While it also allows cells to grow and divide in the complete absence of FtsN, *ftsB*^*E*56*A*^, Δ*ftsN* cells divide less efficiently than wt and are modestly elongated (29) (Fig. 5a, second panel, Supplementary Table 7). In *ftsB*^*E*56*A*^ cells, we observed a significant increase in the slow-moving population of FtsW (Fig. 5b, top panel, orange curves); the fast-moving, FtsZ-dependent population of FtsW-RFP was essentially abolished, and nearly all directional moving FtsW-RFP molecules moved at an average speed of ~9nm/s (9.3±0.8 nm, 93.3±3.7%, N=176 segments, Fig. 5b and c, orange, Supplementary Table 6). Most interestingly and as expected, in *ftsB*^*E*56*A*^, Δ*ftsN* cells where FtsN is absent, the slow-moving population of FtsW-RFP was reduced while the fast-moving population recovered to a level close to that in WT cells (23.8±4.4%, N=144 segments, Fig. 5b second panel, Supplementary Table 6). These results indicate that even though FtsN is no longer essential in the superfission *ftsB*^*E*56*A*^ background, it still contributes to the transitioning of FtsW from the fast-moving, FtsZ-dependent state to the slow-moving, sPG synthesis-dependent mode.

## Discussion

Taken together, our data support a two-track model (Fig. 6), in which directionally moving FtsW and FtsI, and possibly other sPG remodeling enzymes and regulators as well, occupy at least two “tracks” within the septum: a fast “Z-track” representing inactive molecules end-trailing treadmilling FtsZ polymers(22), and a slow “sPG-track” representing active molecules that exited the Z-track to produce sPG. Based on the speed of slow-moving FtsW and FtsI, the sPG polymerization rate in live *E. coli* cells is likely ~6 to 14 disaccharides per second (one disaccharide is ~1 nm (32)), on par with what we estimated from previous biochemical labeling experiments (33) (Supplementary). Stationary FtsWI molecules are likely bound to the middle of FtsZ filaments or trapped at sPG synthesis sites waiting for available lipid II or other factors to initiate/continue sPG synthesis, or to depart the sPG-track to engage FtsZ again. FtsN promotes the release of inactive sPG synthase from treadmilling FtsZ polymers to pursue the sPG-track for active synthesis. In this scenario, FtsWI may associate with regulators of sPG synthesis activity such as FtsN and FtsQLB on either or both tracks, and switch between active and inactive states based on appropriate input by the regulatory proteins. Future SMT studies on the dynamics of the FtsQLB and FtsN proteins should help elucidate how they control FtsWI activity and cell constriction in molecular details. The two-track model integrates spatial information into the regulation of sPG synthesis and suggests a novel mechanism for the spatiotemporal coordination of bacterial cell wall constriction.

**Fig. 6.**
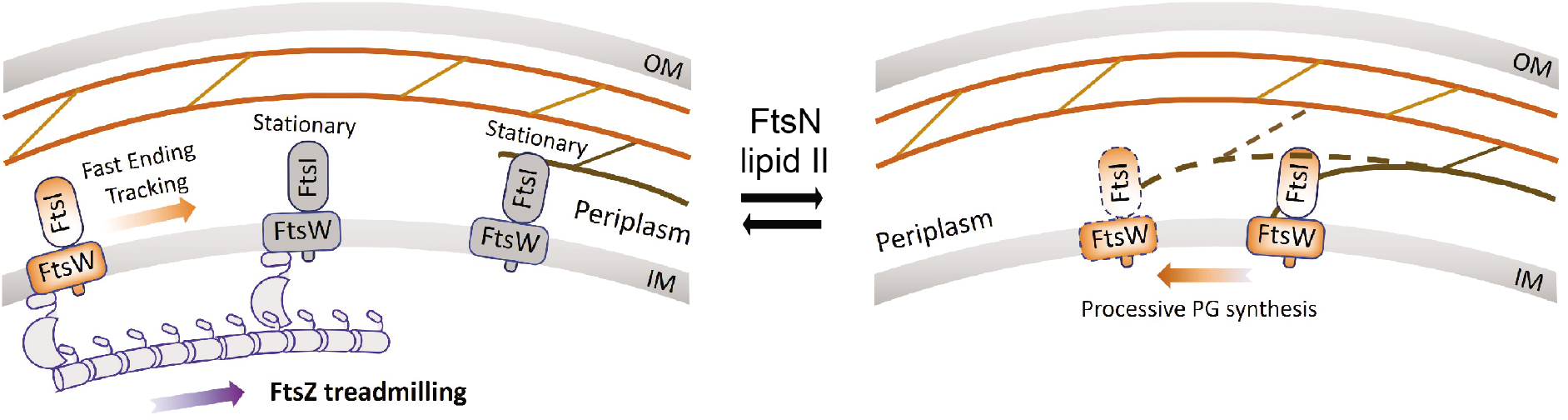
A two-track model integrating spatial information into the regulation of sPG synthase activity. Inactive synthase FtsWI complex follows the treadmilling FtsZ filament (Z-track, left) and is transported to different locations along the septum. Active FtsWI complex engages in processive septal cell wall synthesis along the sPG track (right). FtsN and the available level of cell wall synthesis precursor Lipid II play important roles in promoting the release of inactive FtsWI from the Z-track to pursue sPG synthesis on the sPG-track. Stationary FtsWI complexes (grey molecules, left panel) likely include those bound to internal subunits of FtsZ filaments until the shrinking end approaches, and those bound at sPG synthesis sites waiting for factors (e.g. lipid II substrate or the activator FtsN) to start or continue sPG synthesis.

## Acknowledgements

The authors thank lab members in the Xiao and de Boer labs for helpful discussions and technical assistance, Dr. G. Hauk for sharing plasmids and the CRISPR-Cas9/λ-red recombineering cloning method, Dr. D. S. Weiss for strain EC1908, plasmid pDSW406, anti-FtsN serum, and helpful suggestions on FtsW immunoblotting, Dr. T. Berhardt for strain HC532 and plasmid pHC808, Dr. C. Hale for plasmid pCH650, Dr. E. Goley for help on cell growth measurement, and Dr. R. Tsien for the TagRFP-T construct. This work was supported by NIH GM57059 (to P.d.B), NIH R01 GM086447 (to J.X.), GM125656 (subcontract to J. X.), NSF EAGER Award MCB-1019000 (to J.X.), and a Hamilton Smith Innovative Research Award (to J.X.)

