## Supplemental, Methods, and Extended Data Figures for "FtsW exhibits distinct processive motions driven by either septal cell wall synthesis or FtsZ treadmilling in *E. coli*"

**This PDF file includes:**

*Methods*
*Extended Data Figures 1 to 15*
*Supplementary Table 1 to 8*
*Captions for Movies S1 to S9*
*References*

**Other Supplementary Materials for this manuscript includes the following:**

*Movies S1 to S9*

### *Methods*

#### *Strains, plasmids and growth media:*

Cells were grown in Luria-Bertani (LB) medium, EZ Rich Defined Medium (EZRDM)[1], M9-glucose minimal medium (0.4% D-glucose, 1X MEM amino acids, and 1X MEM Vitamin), or M9-acetate minimal medium (3.4 g/L acetate, pH = 7.0, 1X MEM amino acids, and 1X MEM Vitamin) as specified. For single molecule tracking experiments, vitamins are removed from the M9 medium to minimize background. Where appropriate, antibiotics were included at 25 µg/ml (carbenicillin, kanamycin), 35 µg/ml (chloramphenicol), or 50 µg/ml (ampicillin, spectinomycin), unless specified otherwise.

In most cases, new plasmids were assembled by amplifying appropriate DNA fragments followed by In-fusion cloning (TaKaRa, In-Fusion® HD Cloning Kit). The Quikchange Lightening Kit was used for site-directed mutagenesis (SDM) of plasmid DNA when needed. All primers used for Polymerase Chain Reactions (PCR) or SDM are listed in Supplementary Table 2.

The FtsW-TagRFP-T (FtsW-RFP) fusion protein contains a peptide linker of 15 amino acids (GGGGSPAPAPGGGGS) between the C-terminus of FtsW and N-terminus of TagRFP-T. To generate plasmids encoding this fusion protein, the *ftsW* gene was amplified from the chromosome of *E. coli* strain BW25113 with primers 1 and 2, the *tagrfp-t* gene was amplified from plasmid pJB007[2] with primers 3 and 4, and the vector backbone of plasmid pXY027[3] was amplified with primers 5 and 6. The three DNA fragments were then joined to generate plasmid pXY253. Next, the antibiotic resistance gene on pXY253 was swapped from *cat* to *bla* to yield the plasmid pJL018. To do so, all of pXY253 except for the *cat* gene was amplified with primers 9 and 10, the *bla* gene was amplified from plasmid pKD46 with primers 11 and 12, and the two fragments were joined. The -35 region of the *lacI*$^{Q}$ promoter (GTGCAA) on pJL018 was optimized to TTGACA (*lacI*$^{Q1}$) by SDM with primers 7 and 8, to further suppress the basal expression of FtsW-TagRFP-T to a level that is sufficiently low for single molecule tracking[4]. Further SDM of resulting plasmid pXY349 with primers 13-38 yielded plasmids encoding a single cysteine variant of FtsW-RFP (Supplementary Table 1). The TagRFP-T-FtsI (RFP-FtsI) fusion protein is an N-terminal fusion without any linker. The first methionine of FtsI was removed to minimize the expression of WT FtsI. To generate plasmid pXY388, a linker sequence was first removed from plasmid pJB007 by its amplification and recircularization using primers 55 and 56 by In-fusion cloning. The antibiotic resistance gene was then swapped from *cat* to *aph* and the *lacI*$^{Q}$ was replaced by *lacI*$^{Q1}$, as described above.

Plasmid pCH650 for UppS overexpression was obtained by replacing the 24 bp *Xba*I-*Hind*III fragment of pBAD33 with the 804 bp *Xba*I-*Hind*III fragment of pHC808.

For measuring cell constriction rates by time lapse imaging, plasmid pXY018 was modified to encode mNeonGreen-ZapA, rather than GFP-ZapA, resulting in a brighter and more stable fluorescent fusion protein. The *mNeonGreen* gene was amplified with primers 57 and 58 from the chromosome of strain JXY263, which contains the sandwich *ftsZ-mNeonGreen* fusion gene[5]. The vector backbone containing the *zapA* gene was amplified from plasmid pXY018 with primers 59 and 60. The two DNA fragments were then joined to generate plasmid pXY677.

To generate the FtsW-depletion strain JXY304/pXY287, we first constructed a readily curable plasmid expressing wildtype FtsW under arabinose induction. The *Bacillus subtilis sacB* gene, which confers sensitivity to sucrose in *E. coli*, was amplified from pGH34 (gift from Dr. Glenn Hauk) with primers 39 and 40. The plasmid pDSW406 (pBAD33-*ftsW*) was linearized and amplified using primers 41 and 42. In-fusion joining of the two fragments yielded plasmid pXY287. BW25113 wildtype cells were transformed with pXY287, and the chromosomal *ftsW* gene of BW25113/pXY287 cells was replaced with the *aph* (kanamycin resistance) gene using λ-red recombination[6] with an *aph* fragment that was amplified from plasmid pKD13 using primers 43 and 44. As expected, the resulting strain (JXY304/pXY287) is not viable in the absence of L-arabinose or in the presence of 6% sucrose.

To create a chromosomal *ftsW-tagrfp-t* allele, a fragment encoding the 15-residue linker (see above) and TagRFP-T was inserted immediately downstream of *ftsW* in strain BW25113 using a coupled CRISPR-Cas9/λ-red recombineering system developed by Dr. Glenn Hauk. First, a potential sgRNA-binding DNA sequence near the 3' end of *ftsW* gene (AGGTTCACGATGAGTGGTCA**AGG**) was selected using ChopChop[7]. Primers 45-48 were then used to construct plasmid pJM19, which encodes a sgRNA targeting the original AGGTTCACGATGAGTGGTCA sequence. Briefly, the sgRNA sequence was created using primers 47 (which includes the protospacer region) and 48, with pGH34 as template, and the fragment was inserted in pGH34 that had been amplified with primer 45 and 46. Next, a dsDNA fragment bearing *linker-tagrfp-t* flanked by 50 bp sequences homologous to the *ftsW* locus, was amplified from pXY349 with primers 51 and 52. BW25113 cells were then co-transformed with plasmids pKD46 and pGH33, and a 50 ml culture of BW25113/pKD46/pGH33 cells was grown at 30ºC to log-phase ($OD_{600}$ = 0.5). Arabinose was added to a final concertation of 0.2%. After another hour of growth, cells were harvested and used to prepare a 0.5 ml suspension of electrocompetent cells using a standard protocol (Short Protocols in Molecular Biology, Chapter 1). An aliquot (75 µl) was then used for electro-transformation with 100 ng pJM19 and 500 ng of the *linker-tagrfp-t* fragment. After recovery at 30ºC for 90 mins, the entire mixture was plated on LB agar containing 60 µg/ml carbenicillin (CB), 50 µg/ml kanamycin (KAN), and 75 µg/ml chloramphenicol (CAM), and incubated at 30ºC for 30 hrs. The CRISPR-Cas9 complex in these cells was designed to cut the native *ftsW* gene, causing most cells to die. However, cells bearing the desired *ftsW-tagrfp-t* gene should survive, as this recombinant allele is not recognized by the sgRNA. Survivors were screened by colony-PCR and cured of the three plasmids by growth at 37ºC in the presence of 6% sucrose and absence of antibiotics. Presence of the chromosomal *ftsW-tagrfp-t* allele in the cured strain (JXY422) was verified by colony-PCR again. Strains with the chromosomal *ftsW*$^{I302C}$ allele (JXY559, JXY564) or *murJ*$^{A29C}$ allele (JXY589) were generated using a similar method. Plasmids pXY550 and pXY588 encoding the sgRNAs targeting wild type *ftsW* or *murJ* gene were obtained by the same method as pJM19, but by using primers 49 (*ftsW*) and 50 (*murJ*) instead of 47. In this case, λ-red recombineering was performed with a mutagenic ssDNA fragment harboring the appropriate single codon mutation and flanking sequences (primer 53 and 54 for *ftsW*$^{I302C}$ and *murJ*$^{A29C}$ respectively).

To track FtsW-TagRFP-T molecules in *ftsZ* mutant cells, JXY001 (*ftsZ*$^{G105S}$), JXY002 (*ftsZ*$^{E250A}$), JXY003 (*ftsZ*$^{D158A}$), JXY005 (*ftsZ*$^{E238A}$), and JXY006 (*ftsZ*$^{D269A}$) were transformed with plasmid pXY349.

The isolation and characterization of the *ftsI*$^{R167S}$ and *ftsW*$^{E289G}$ superfission alleles will be described in extensive detail in a future publication elsewhere. Briefly, both alleles were selected from plasmid libraries with randomly mutagenized *ftsW* or *ftsI*, based on the abilities of the library plasmids to i) confer viability to Δ*ftsN* cells, and ii) to suppress the division defects of strain TB77 (*ftsN*$^{slm117}$). The *ftsI*$^{R167S}$ and *ftsW*$^{E289G}$ alleles were next moved onto a *repA*$^{ts}$ pSC101-derivative to allow for allelic exchange between the resulting plasmids and the chromosome by the method of Hamilton et al[8]. Such exchange in cells of strain TB77 then resulted in strains PM4 (*ftsN*$^{slm117}$ *ftsI*$^{R167S}$) and PM15 (*ftsN*$^{slm117}$ *ftsW*$^{E289G}$), respectively. Strain PM6 was next obtained by P1-

mediated co-transduction of *ftsI*$^{R167S}$ with *leu*+ from PM4 to BL17, and PM8 by transduction of *ftsN*::*aph* (Δ*ftsN*) from CH34/pCH201 to PM6. Similarly, strain PM17 was obtained by co-transduction of *ftsW*$^{E289G}$ with *leu*+ from PM15 to BL17, and PM18 by transduction of *ftsN*::*aph* from CH34/pCH201 to PM17.

#### *Analyses of FtsW-RFP single cysteine mutants*

To test single cysteine variants of FtsW-RFP for inactivation with MTSES, plasmids listed in Supplementary Table 1 were introduced into strain JXY304/pXY287. Purified transformants were grown overnight at 37$^{o}$C in LB medium containing 0.2% L-arabinose. Cultures were then diluted 1000 times in M9 medium with 0.4% D-glucose and 100 µM IPTG to simultaneously suppress expression of native FtsW from pXY287 and induce expression of the single cysteine FtsW-RFP variant from the newly introduced plasmid. Cultures were split in two and incubated at 37$^{o}$C until they reached a cell density between 0.2-0.3 ($OD_{600}$). MTSES (to 1 mM) was added to one of each duplicate culture, and growth was continued for 3-4 additional cell cycles (~ 3 hrs). Cells from 1 mL of each culture were fixed with 3.8% paraformaldehyde for 20min at RT, and then imaged by bright field microscopy. Observed cell morphologies were categorized into four classes (Extended Data Fig 1): 1. Near normal morphology, regardless of MTSES treatment. 2. MTSES causes increased cell length, but not the formation of long, septated cell chains. 3. MTSES causes the formation of long cell chains with partially completed septa. 4. Formation of long cell chains or filaments, even in the absence of MTSES.

#### *Western blotting*

For FtsW-TagRFP-T immunoblotting, 25 mL BW25113 and JXY422 cells were harvested by centrifugation at $OD_{600}$ ~ 0.3 and resuspended in 100 µL 1X BugBuster Protein Extraction Reagent (MilliporeSigma). The mixture was constantly agitated in a rotator at RT for 3 – 4 hours. 100 µL 2X SDS gel-loading buffer was then add to the tube and mix by pipetting. The mixture was agitated in a rotator for another 3 hours at RT or incubated at 4 ºC overnight. 20uL of the mixtures were applied to a 10% SDS-PAGE gel and electroblotted onto nitrocellulose membranes. Note that because FtsW is prone to aggregation, the whole process was performed at RT or 4 ºC. After blocking in TBS with 0.05% (v/v) Tween-20 and 5% (w/v) non-fat dry milk, membranes were incubated with RFP Polyclonal Antibody (ThermoFisher, R10367) diluted 1:2000 in TBS with 0.05% (v/v) Tween-20 and 1% (w/v) BSA. FtsW-TagRFP-T bands were detected using goat anti-rabbit HRP (1:50,000 dilution; Bio-Rad) and Clarity TM Western ECL Substrate (Bio-Rad). Band intensities were quantified using ImageJ[9] Gel plugin.

For FtsN immunoblotting, 3mL MG1655 and EC1908 (with or without Arabinose) cells were harvested by centrifugation at $OD_{600}$ ~ 0.3 and resuspended in 50 µL M9 medium. 50 µL 2X SDS gel-loading buffer was then add to the tube and boiled at 95 ºC for 10 min. 10 uL of the mixtures were applied to a 10% SDS-PAGE gel and electroblotted onto nitrocellulose membranes. After blocking in TBS with 0.05% (v/v) Tween-20 and 5% (w/v) non-fat dry milk, membranes were incubated with anti-FtsN serum (a gift from Dr. David Weiss) diluted 1:5000 in TBS with 0.05% (v/v) Tween-20 and 1% (w/v) BSA. FtsN bands were detected using goat anti-rabbit HRP (1:50,000 dilution; Bio-Rad) and imaged after treatment with Clarity TM Western ECL Substrate (Bio-Rad). Band intensities were quantified using ImageJ plugin.

***Glycan strain labeling using NAM-KU system***

Cultures of the appropriate strains harboring plasmid pBBR1-KU were grown overnight at 37 ºC in LB medium, diluted 1:200 into 3 ml of M9 medium, and grown further at 25 ºC to a cell density of ~0.6 ($OD_{600}$). Fosfomycin (to 100 μg/ml) was added to suppress endogenous production of UDP-NAM, and incubation was continued for 30 min. At this point, IPTG (to 1 mM) was added to induce production of the plasmid-encoded AmgK and MurU enzymes and, when indicated, MTSES (to 1 mM) was added to inactivate the relevant single-cysteine variant of MurJ, PBP1B and/or FtsW. After incubation for another 30 min, cells were pelleted at 4110 rpm for 10 min at 25 ºC and resuspended in 200 μL of the supernatant. Alkyl-NAM (to 2 mg/ml) was then added and, after incubation for another 30 min, cells were fixed and permeabilized using 70% ethanol as previously described[2]. After fixation, cells were washed three times in PBS and fluorescently labeled using the Click-iT™ Plus Alexa Fluor™ 647 Pixolyl Azide Toolkit (Invitrogen, No. C10643).

Cells were imaged on an Olympus IX-71 inverted microscope equipped with a 1.4 NA 100X phase objective, 1.6X magnification enabled, and an Andor iXon-DU897 camera. Every field of view was imaged with five Z-stacks from the bottom of the cell in 200 nm increments. The fluorophores were excited by 2 mW 647 nm excitation laser and detected with ET700/75 emission filter (Chroma) using a 100ms exposure time.

The Z-stack images were first summed up for all Z planes. The background of the image was then subtracted out using the outside region. The cell outline and a ~250 nm (3 pixels) wide mid-cell band were segmented out using the phase images, where the average fluorescence intensity in the septum ($I_r$) and the whole cell ($I_c$) were calculated. The cells with observable septal labeling were manually picked and counted as cells with sPG synthesis ($N_{sep}$) (Fig 1b). The total sPG synthesis in those cells was quantified as: $I_{sep} = (I_r - I_c) \times A_r$, where the $A_r$ is the area of the septa. The overall loss of the sPG synthesis is calculated as:

$$L_{sPG} = (1 - \frac{P_{r,MTSES}}{P_{r,no\ MTSES}} \times \frac{I_{sep,MTSES}}{I_{sep,no\ MTSES}})\ 100\%$$

Where the $P_r$ is the percentage of cells with septal labeling ($N_{sep}/N_{cell}$), and MTSES indicates the presence of MTSES treatment.

#### *Cell growth and constriction rate measurement*

The cell growth rate in different growth media was measured by monitoring the $OD_{600}$ of shaking cultures every 30 min (EZRDM), 1 hour (M9-glucose and M9-acetate), and fitting the exponential part of the growth curve.

The constriction rate was measured using time-lapse images of mNeonGreen-ZapA as a fluorescent marker of the septum. PM6/pXY677 cells were grown in different growth media (M9-glucose, M9-acetate, EZRDM) to log phase and placed on corresponding agarose gel-pads. The cells were imaged every 1 minute (M9-glucose or EZRDM) or 3 minutes (M9-acetate) for 30 frames on an Olympus IX-71 inverted microscope equipped with a 1.49 NA 100X objective and an Andor DU897 camera in epifluorescence-illumination mode using a 488-nm laser (Coherent Obis). The laser power was around $0.1 W/cm^2$, without noticeable photo-toxicity on cell growth rate.

The fluorescent images were first aligned to correct mechanical drift using an ImageJ plugin StackReg[10]. The images were then smoothed by a simple moving average method (n = 3 frames) to make kymographs (Extended Data Fig. 14b) of the septum using ImageJ Kymograph plugin (written by J. Rietdorf and A. Seitz). The constriction rate was estimated from the sum of the slope of the edges on both sides in the kymograph.

***Single molecule tracking of FtsW-RFP and data analysis***

FtsW-RFP tracking was performed on the similar optical set up as previously described (an Olympus IX-81 inverted microscope equipped with a 1.45 NA 100X objective and an Andor DU888 camera in epifluorescence-illumination mode using a 561-nm laser (Coherent Sapphire)[2]. The focal plane was placed at ~200nm from the bottom of the cell to image the molecules moving on the bottom half of the cylindrical portion of the cell body. Single FtsW-RFP molecules were tracked continuously with 0.5 second exposure time for 200 s using a laser power of 2 W/cm$^2$.

Before imaging, cells were grown to log phase ($OD_{600}$ = 0.1 – 0.3) at 25°C without IPTG induction in M9-glucose or M9-acetate medium. In EZRDM, cells in saturated culture were reinoculated 1:100 to fresh medium with 10 µM IPTG (and 0.2% L-arabinose for UppS induction if needed) and allowed to grow at 25°C for 3 hours to reach log phase. Cells were then loaded onto a 50 μL, 3% agarose gel pad (containing the same growth medium). For drug-treated conditions, 0.5 μL appropriate drug solution was added on top of the gel pad right before applying cells. The moment of the latter was counted as time zero and all images were collected within three hours of drug treatment (1.5 hours for Aztreonam treatment). The final concentrations used were: 1 µg/ml Aztreonam, 200 µg/ml Fosfomycin, and 0.1 mM MTSES. For the fixed-cell control, log phase cells were fixed as described previously[3] and placed on a gel pad containing phosphate buffered saline.

Single FtsW molecules were first localized using 2D gaussian fitting in an ImageJ[9] plug-in, ThunderSTORM[11]. A bandpass filter (70 – 400nm) for sigma was applied to remove the single pixel noise and out-of-focus molecules. The resulting localizations were linked to trajectories using a nearest neighbor algorithm[12]. The distance threshold was set to 300 nm/frame, which approximates a maximum D ~ 0.05 µm$^2$/s or a maximum speed of 600 nm/s. To link blinking localizations from the same molecules, a time threshold of 5 frames was chosen according to the off-time distribution. Only trajectories near the midpoint of the cell's long axis or near visible constriction sites were used in the analysis.

One bias that arises when tracking molecules on the rod-shape cell envelope is that the real displacement around the circumference is underestimated (Extended Data Fig. 5). If the cell diameter or radius is known, the real position of a molecule on the cell envelope can be back-calculated. Therefore, we grew strain BW25113/pJL005 in M9-glucose, and for each of 55 cells estimated its diameter using its bright field image. We first line-scanned the bright field image intensity across the cell division site and determined a bright field cell diameter between the cross points generated from the average background and the edge of the central part in the cell (Extended Data Fig. 5). We then applied 3D super-resolution imaging on the Z-ring from each of the same cells, as previously described[13]. The Z-ring image was then fit to a circle to estimate the Z-ring diameter. On average, the bright field diameter was found to be 57 nm greater than the Z ring diameter. Considering a ~17nm distance from the inner-membrane to the Z ring[3], we can estimate that the ‘real’ cell diameter is 23 nm smaller than the bright field measurement. The ‘real’ position of a molecule on the cell circumference is then calculated as shown in Extended Data Fig. 5c. Note that the position along the long axis of the cell does NOT change due to the cylindrical geometry of the cell.

Unwrapped, corrected trajectories as described above were then segmented to determine whether a FtsW-RFP molecule in a segment is stationary or moving directly. To classify directional movement and minimize bias, we first fit the segment with a line. Then we defined a metrics $R$, which is the ratio of the displacement and the standard deviation of fitting residuals ($S$) (Extended Data Fig. 5e). Segments from directionally moving molecules would produce smaller

$R$ values than the stationary ones considering similar diffusion coefficients and localization error from imaging. Therefore, we constructed the probability density function of $R$ by simulating trajectories at a specific directional speed with the typical diffusion coefficient, confinement, and localization error obtained from our experiment. For individual segments, we can infer the probability of the molecule undergoing directional movement ($P_m$, see the **Segment Classification Section** below for simulation and derivation details). In the paper, we classify a segment as directional with an empirical threshold of $P_m$ and $R$. All the other segments are stationary if the $S$ is smaller than 50 nm (confined in a ~100nm range). Since the confidence of classification is correlated with the segmentation length, we only consider segments longer than 6 frames to minimize classification error. We note that the velocity estimated from MSD curve fitting is not accurate when the dwell time of the segment is short (< 20 frames) or when there is more than one moving state in a single trajectory (Fig 2b, Extended Data Fig. 6).

To estimate the percentage of time FtsW-TagRFP-t spends in directional moving states, the duration of time of all directional moving segments were summed and divided by the total time of all trajectories. The error was estimated by bootstrapping 1000 times. The ensemble average speed was also estimated by bootstrapping 1000 times of all the speeds.

The cumulative probability density functions (CDF) of the direction moving speeds were calculated for each condition and fit by either single (for FtsZ treadmilling) or double lognormal populations for FtsW (or FtsI) tracking:

$$CDF = P_1 \cdot \frac{\left(1 + \operatorname{erf}\left[\frac{\ln v - \mu_1}{\sqrt{2}\sigma_1}\right]\right)}{2} + (1 - P_1) \cdot \frac{\left(1 + \operatorname{erf}\left[\frac{\ln v - \mu_2}{\sqrt{2}\sigma_2}\right]\right)}{2}$$

Where the $v$ is the FtsW (or FtsI) moving speed and $P_1$ is the percentage of the first population. For a single population model, $P_1 = 1$. Two parameters $\mu$ and $\sigma$ are the natural logarithmic mean and standard deviation.

The average speed of each population is calculated as: $\exp\left(\mu + \frac{\sigma^2}{2}\right)$. To estimate the error of the speed and percentage (Supplementary Table 3-6), the CDF curves were bootstrapped 1000 times and fit with the corresponding equation (single- or double-population).

***Segment Classification***

As shown in Extended Data Fig. 5e, we can linearly fit a single molecule trajectory (segment) with $T$ frames to estimate the speed $V$ and calculate the standard deviation of fitting residuals $S$, while the ground truth of the speed is $\hat{V}$. The confidence of calling one segment as in directional motion depends on the related level of $S$, which is a function of the localization error, diffusion coefficient and confinement size. We further defined a dimensionless metric $R$ that is $S$ normalized by the total displacement of the segment $L = T \cdot V$. Thus, the classification problem can be described as estimating the probability of the segment to be 'stationary' given $T$ and $R$:

$$P_s = P(Stationary|T,R) \approx P\left(\hat{V} = 0 \middle| T,R\right) \tag{1}$$

And the probability of directional movement is

$$P_m = P(processive|T,R) \approx P\left(\hat{V} = V \middle| T,R\right) \tag{2}$$

Here, we simplified the problem to a binary classification with two most likely speeds, 0 or $V$.

We developed the following algorithm to calculate the probabilities in (1) and (2) and to classify a certain segment as stationary or directional moving.

**Step 1: build the probability density function (PDF) of $R$ from simulation**

Since $R$ is defined as $\frac{S}{T \cdot V}$, where S depends on the localization error ($\delta$), diffusion coefficient ($D$) and confinement size ($B$), we first determined $\delta_0 = 37nm, D_0 = 0.0007$ μm$^2$/s, $B_0 = 95\ nm$ of FtsW-RFP under our experimental condition from the MSD fitting results shown in Extended Data Fig. 7c. Note that we simulated stationary RFP-FtsI molecules' PDF using the same parameter set since FtsI is expected to be in the same complex and displays similar motion as FtsW-RFP. For an experimental segment with length $T_{exp}$ and fit speed $V_{fit}$, we carried out two sets of random simulations to generate 1000 segments based on fixed $\delta_0, D_0,\ B_0,\ T_{exp}$, and $V = 0\ or\ V_{fit}$. All the simulated segments were then fit linearly to calculate their $R$ values. The distribution was used to construct the PDF of $R$ given a stationary or directional movement state:

$$PDF_s(R) = PDF\left(R \middle| \delta_0, D_0, B_0, T_{exp}, V = 0\right) \quad (3)$$

$$PDF_m(R) = PDF\left(R \middle| \delta_0, D_0, B_0, T_{exp}, V = V_{fit}\right) \quad (4)$$

**Step 2: Estimate the experimental distribution of $R_{exp}$:**

We determined the distribution of $R_{exp}$ using a bootstrapping-like procedure. We first randomly deleted 10% of the datapoints from the segment for 100 times. We then fit all the re-sampled segments by line and calculated the corresponding standard error ($\sigma$) of $R_{exp}$. Next, we integrated $\langle R_{exp} \rangle \pm \sigma$ in (3) and (4) to obtain the probability of $R_{exp}$ given the conditions described above and the speed.

$$P\left(R_{exp} \middle| stationary\right) = \int_{\langle R_{exp} \rangle - \sigma}^{\langle R_{exp} \rangle + \sigma} P\left(R \middle| \delta_0, D_0, B_0, T_{exp}, V = 0\right) dR \quad (5)$$

$$P\left(R_{exp} \middle| processive\right) = \int_{\langle R_{exp} \rangle - \sigma}^{\langle R_{exp} \rangle + \sigma} P\left(R \middle| \delta_0, D_0, B_0, T_{exp}, V = V_{fit}\right) dR \quad (6)$$

**Step 3: Bayesian inference:**

For every segment, we calculated its $P_m$ using Bayes' theorem:

$$P_m = \frac{P(R_{exp}|processive)P(processive)}{P(R_{exp}|stationary)P(Stationary) + P(R_{exp}|processive)P(processive)}$$

$$= \frac{P(R_{exp}|processive)}{P(R_{exp}|stationary) + P(R_{exp}|processive)}$$

Here we assume no prior knowledge on the type of motion, thus $P(stationary) = P(processive)$ .

**Step 4: Classification:**

We used a set of thresholds: $P_m > 0.75$ and $R_{exp} < 0.4$ to identify directional or processive moving molecules, which is comparable with a manual classification but largely reduced the potential systematic bias.

***Estimation of PG synthesis rate in E. coli***

Burman et al.[14] estimated that *E. coli* cells growing with a doubling time of 48 min, insert about 450,000 new muropeptides in the peptidoglycan sacculus during an 8-min labeling period with radioactive cell wall precursors. This is equivalent to 150 PG loops around the cell circumference (each loop corresponds to ~ 3000 muropeptides or 3000 nm). If one assumes that there are ~150 PG synthase complexes[15] that simultaneously synthesize the PG in a processive fashion, the polymerization rate for each loop would be 3000 nm / 480 s = 6 nm/s. Note that this number is lower than the directional moving speed of the RodA/PBP2 system for cell wall elongation[16] , and that this simplified estimation does not take into account the number of enzyme molecules presented on each loop's synthesis, the contribution of other PGTases, the growth condition (our growth condition is at RT and in minimal media), and the simultaneous PG turnover process. Nevertheless, the speed estimated from these labeling experiments and the speed of the RodA system agree with our SMT measurements in the same order of magnitude.

Supplementary Table 1: Strains and plasmids used in the study

| **Strain** | **Genotype** | **Reference/source** |
|---|---|---|
| BW25113 | *Δ(araD-araB)567, ΔlacZ4787(::rrnB-3), rph-1, Δ(rhaD-rhaB)568, hsdR514* | 17 |
| BL17 | TB28, *leu*::Tn10 | 18 |
| BL167 | TB28 *ftsB*$^{E56A}$ | 18 |
| BL173 | TB28 *ftsB*$^{E56A}$ *ftsN*::*aph* | 18 |
| CH34* | TB28 *ftsN*::*aph* | 19 |
| EC1908* | MG1655 $P_{BAD}$::*ftsN* | 20 |
| HC532 | MG1655 *lysA::frt ponA::frt pbpC::frt mtgA::frt ponB*$^{S247C}$ | 16 |
| JXY001 | BW25113 *ftsZ*$^{G105S}$(ts) | 2 |
| JXY002 | BW25113 *ftsZ*$^{E250A}$ | 2 |
| JXY003 | BW25113 *ftsZ*$^{D158A}$ | 2 |
| JXY005 | BW25113 *ftsZ*$^{E238A}$ | 2 |
| JXY006 | BW25113 *ftsZ*$^{D269A}$ | 2 |
| JXY263 | BW27783 *ftsZ*::*ftsZ*55-56-*mNeonGreen*, $P_{lac}$::*ftsZ-tagRFP-t* | 2 |
| JXY304* | BW25113 *ftsW*::*aph* | This study |
| JXY422 | BW25113 *ftsW-tagrfp-t* | This study |
| JXY559 | BW25113 *ftsW*$^{I302C}$ | This study |
| JXY564 | MG1655 *lysA::frt ponA::frt pbpC::frt mtgA::frt ponB*$^{S247C}$ *ftsW*$^{I302C}$ | This study |
| JXY589 | BW25113 *murJ*$^{A29C}$ | This study |
| MG1655 | *ilvG rfb*50 *rph*1 | 21 |
| PM4 | TB28 *ftsN*$^{slm117}$ *ftsI*$^{R167S}$ | This study |
| PM6 | TB28 *ftsI*$^{R167S}$ | This study |
| PM8* | TB28 *ftsI*$^{R167S}$ *ftsN*::*aph* | This study |
| PM17 | TB28 *ftsW*$^{E289G}$ | This study |
| PM18 | TB28 *ftsW*$^{E289G}$ *ftsN*::*aph* | This study |
| TB28 | MG1655 *lacIZYA*::*frt* | 22 |
| TB77 | TB28 *ftsN*$^{slm117}$ (*ftsN*::EZTnKan-2) | 19 |
| **Plasmid** | **Genotype** | **Reference/source** |
| pAD004 | ColE1, *bla lacI*$^{Q1}$ $P_{T5lac}$::*ftsW*$^{I302C}$*-tagRFP-t* | This study |
| pAD005 | ColE1, *bla lacI*$^{Q1}$ $P_{T5lac}$::*ftsW*$^{I303C}$*-tagRFP-t* | This study |
| pBAD33 | pACYC, *cat araC* $P_{BAD}$:: | 23 |
| pBBR1-KU | ColE1, *cat* $P_{lac}$::*amgK murU* (*P. putida* genes) | 24 |

| | | |
|---|---|---|
| pBL145 | pSC101, *aadA cI*$^{ts}$ $P_{\lambda R}$::*ftsN*$^{1-90}$ | 18 |
| pCH201 | ColE1, *bla* $^{Q}$*lacI*$^{Q}$ $P_{lac}$::*gfp-ftsN* | 25 |
| pCH650 | pACYC, *cat araC* $P_{BAD}$::*uppS* | This study |
| pDSW406 | pACYC, *cat araC* $P_{BAD}$::*ftsW* | 26 |
| pGH33 | ColE1, *aph* $P_{BAD}$::*spCas9* | A gift from Dr. Glenn Hauk |
| pGH34 | pACYC, *cat* $P_{BAD}$::*sacB, sgRNA* | A gift from Dr. Glenn Hauk |
| pHC808 | ColE1, *cat lacI*$^{Q}$ $P_{tac}$::*uppS* | 27 |
| pJB007 | ColE1, *cat lacI*$^{Q}$ $P_{T5lac}$::*tagrfp-t-ftsI* | 2 |
| pJL005 | ColE1, *bla lacI*$^{Q}$ $P_{T5lac}$::*ftsZ-meos3.2* | 13 |
| pJL018 | ColE1, *bla lacI*$^{Q}$ $P_{T5lac}$:: *ftsW-tagRFP-t* | This study |
| pJM19 | pACYC, *cat* $P_{BAD}$::*sacB, ftsW-sgRNA* | This study |
| pKD13 | R6K, *bla frt::aph::frt* | 17 |
| pKD46 | pSC101, *bla repA*$^{ts}$, *araC* $P_{BAD}$::*gam bet exo* | 17 |
| pXY018 | ColEl, *cat lacI*$^{Q}$ $P_{T5lac}$::*gfp-zapA* | 2 |
| pXY027 | ColE1, *cat lacI*$^{Q}$ $P_{T5lac}$::*ftsZ-gfp* | 3 |
| pXY253 | ColE1, *cat lacI*$^{Q}$ $P_{T5lac}$::*ftsW-tagRFP-t* | This study |
| pXY287 | pACYC, *cat araC* $P_{BAD}$::*ftsW, sacB* | This study |
| pXY349 | ColE1, *bla lacI*$^{Q1}$ $P_{T5lac}$::*ftsW-tagRFP-t* | This study |
| pXY388 | ColE1, *aph lacI*$^{Q1}$ $P_{T5lac}$:: *tagRFP-t-ftsI* | This study |
| pXY437 | ColE1, *bla lacI*$^{Q1}$ $P_{T5lac}$::*ftsW*$^{A301C}$*-tagRFP-t* | This Study |
| pXY438 | ColE1, *bla lacI*$^{Q1}$ $P_{T5lac}$::*ftsW*$^{L367C}$*-tagRFP-t* | This study |
| pXY439 | ColE1, *bla lacI*$^{Q1}$ $P_{T5lac}$::*ftsW*$^{E289G}$*-tagRFP-t* | This study |
| pXY441 | ColE1, *bla lacI*$^{Q1}$ $P_{T5lac}$::*ftsW*$^{S131C}$*-tagRFP-t* | This study |
| pXY442 | ColE1, *bla lacI*$^{Q1}$ $P_{T5lac}$::*ftsW*$^{S136C}$*-tagRFP-t* | This study |
| pXY443 | ColE1, *bla lacI*$^{Q1}$ $P_{T5lac}$::*ftsW*$^{L198C}$*-tagRFP-t* | This study |
| pXY445 | ColE1, *bla lacI*$^{Q1}$ $P_{T5lac}$::*ftsW*$^{L268A}$*-tagRFP-t* | This study |
| pXY446 | ColE1, *bla lacI*$^{Q1}$ $P_{T5lac}$::*ftsW*$^{L276C}$*-tagRFP-t* | This study |
| pXY447 | ColE1, *bla lacI*$^{Q1}$ $P_{T5lac}$::*ftsW*$^{L288C}$*-tagRFP-t* | This study |
| pXY448 | ColE1, *bla lacI*$^{Q1}$ $P_{T5lac}$::*ftsW*$^{A294C}$*-tagRFP-t* | This study |
| pXY449 | ColE1, *bla lacI*$^{Q1}$ $P_{T5lac}$::*ftsW*$^{L307C}$*-tagRFP-t* | This study |
| pXY550 | pACYC, *cat* $P_{BAD}$::*sacB, ftsWI302-sgRNA* | This study |
| pXY588 | pACYC, *cat* $P_{BAD}$::*sacB, murJA29C-sgRNA* | This study |
| pXY677 | ColE1, *cat lacI*$^{Q}$ $P_{T5lac}$::*mNeonGreen-zapA* | This study |

* Cells require a complementing plasmid or specific growth conditions for survival.

Supplementary Table 2: Primers used in the study

| Primer | Sequence | Purpose |
|---|---|---|
| 1 | GAGGAGAAATTAACTACTAGTATGGTGTCTAAGGGCGAAGAGC | Amplification of *ftsW* gene for pXY349 |
| 2 | CGGTGCCGGTGCCGGGCTACCACCGCCACCCTTGTACAGCTCGTCCATGCCA | |
| 3 | CCGGCACCGGCACCGGGCGGTGGCGGTAGCCGTTTATCTCTCCCTCGCC | Amplification of *tagrfp-t* gene for pXY349 |
| 4 | GGTCGACCCTTAGCGGCCGCTTATCGTGAACCTCGTACAAACG | |
| 5 | ACTAGTAGTTAATTTCTCCTCTTTAATG | Amplification of the vector backbone for pXY349 |
| 6 | GCGGCCGCTAAGGGTCG | |
| 7 | TGCCATACCGCGAAAGGTTTGTCAACATTCGATGGTGTCGGAATT | Mutagenesis of *lacI*$^{Q}$ promotor for pXY349 |
| 8 | AATTCCGACACCATCGAATGTTGACAAACCTTTCGCGGTATGGCA | |
| 9 | TTTTTTTAAGGCAGTTATTGGTGC | Amplification of the vector to swap *cat* to *bla* |
| 10 | ACGTCTCATTTTCGCCAAAAGTT | |
| 11 | GCGAAAATGAGACGTTTACCAATGCTTAATCAGTGAGG | Amplification of *bla* gene |
| 12 | ACTGCCTTAAAAAAACGCGGAACCCCTATTTGTTTA | |
| 13 | ACCCCAGTTCTTCGCCGATACAGGCGAAAATAAAGTCAGTG | Mutagenesis of *ftsW* for pAD004 |
| 14 | CACTGACTTTATTTTCGCCTGTATCGGCGAAGAACTGGGGT | |
| 15 | CCAGTTCTTCGCCGCAAATGGCGAAAATAAAGTCAGTGTG | Mutagenesis of *ftsW* for pAD005 |
| 16 | CACACTGACTTTATTTTCGCCATTTGCGGCGAAGAACTGG | |
| 17 | AGTTCTTCGCCGATAATGCAGAAAATAAAGTCAGTGTGCGC | Mutagenesis of *ftsW* for pXY437 |
| 18 | GCGCACACTGACTTTATTTTCTGCATTATCGGCGAAGAACT | |
| 19 | CTTTGGTCGGGCACATCCCCGCCGCCGCG | Mutagenesis of *ftsW* for pXY438 |
| 20 | CGCGGCGGCGGGGATGTGCCCGACCAAAG | |
| 21 | CGCTTCCGGCAGATACCCCAGTTTTTGTACCGA | Mutagenesis of *ftsW* for pXY439 |
| 22 | TCGGTACAAAAACTGGGGTATCTGCCGGAAGCG | |
| 23 | GCGATGCCCCTTTAACGCAGCTACCCACTACCAG | Mutagenesis of *ftsW* for pXY441 |
| 24 | CTGGTAGTGGGTAGCTGCGTTAAAGGGGCATCGC | |
| 25 | GAGATCGATCCAACGGCATGCCCCTTTAACCGAGC | Mutagenesis of *ftsW* for pXY442 |
| 26 | GCTCGGTTAAAGGGGCATGCCGTTGGATCGATCTC | |
| 27 | ACCACCACCGTACCACAGTCTGGCTGTGCCAG | Mutagenesis of *ftsW* for pXY443 |
| 28 | CTGGCACAGCCAGACTGTGGTACGGTGGTGGT | |
| 29 | CGCCGCGACCAAACGCCATGCACGATTGCGTTAACTGATAG | Mutagenesis of *ftsW* for pXY445 |
| 30 | CTATCAGTTAACGCAATCGTGCATGGCGTTTGGTCGCGGCG | |
| 31 | CTTGCCCCCAACATTCGCCGCGACCAAACGCCA | Mutagenesis of *ftsW* for pXY446 |

| | | |
|---|---|---|
| 32 | TGGCGTTTGGTCGCGGCGAATGTTGGGGGCAAG | |
| 33 | GCGCTTCCGGCAGATACTCGCATTTTTGTACCGAGTTACCTAAAC | Mutagenesis of *ftsW* for pXY447 |
| 34 | GTTTAGGTAACTCGGTACAAAAATGCGAGTATCTGCCGGAAGCGC | |
| 35 | AAAACTGGAGTATCTGCCGGAATGCCACACTGACTTTATTTTCGCC | Mutagenesis of *ftsW* for pXY448 |
| 36 | GGCGAAAATAAAGTCAGTGTGGCATTCCGGCAGATACTCCAGTTTT | |
| 37 | CCACACCGACATACCCGCATTCTTCGCCGATAATGGCGAAAATAA | Mutagenesis of *ftsW* for pXY449 |
| 38 | TTATTTTCGCCATTATCGGCGAAGAATGCGGGTATGTCGGTGTGG | |
| 39 | TTGGCTGTTTTGGCGTTTGTTTAACTTTAAGAAGGAGAATGAAC | Amplification of *sacB* gene for pXY287 |
| 40 | CCGCTTCTGCGTTCTTTATTTGTTAACTGTTAATTGTCCTTGTT | |
| 41 | CGCCAAAACAGCCAAGCTTG | Amplification of the vector backbone for pXY287 |
| 42 | AGAACGCAGAAGCGGTCTGA | |
| 43 | TTGAACAACGAGGCAATGAGTTTGCCCGTCTGGCGAAGGAGTTAGGTTGAGGAGCTGCTTCGAAGTTCCT | Knockout of *ftsW* |
| 44 | CAATCCACCGGTTCCGCCTGCCATCACCATTAATCGCTTTCCTTGACCACTAGCTGCTTCGAAGTTCCT | |
| 45 | GTGCACATTATACGAGCCGATGA | Amplification of the vector backbone for pJM19 |
| 46 | ATCATGGCGACCACACCCGTCCTGTGGATCCTCTAC | |
| 47 | TCGTATAATGTGCACAGGTTCACGATGAGTGGTCAGTTTTAGAGCTAGAAATAGCAAGTTAA | Amplification of the sgRNA for pJM19 |
| 48 | CGGCGTAGAGGATCCACAGGACGGGTGTGGTCGC | |
| 49 | TCGTATAATGTGCACCCATTATCGGCGAAGAACTGGTTTTAGAGCTAGAAATAGCAAGTTAA | To generate the sgRNA region of pXY550 |
| 50 | TCGTATAATGTGCACGCAATTGTCGCCAGAATCTTGTTTTAGAGCTAGAAATAGCAAGTTAA | To generate the sgRNA region of pXY588 |
| 51 | TGCTGTTGCGTATTGATTATGAAACGCGTCTGGAGAAAGCGCAGGCGTTTGTACGAGGTTCACGAGGTGG | Insertion of *tagrfp-t* gene into chromosomal *ftsW locus* |
| 52 | CCACCGGTTCCGCCTGCCATCACCATTAATCGCTTTCCTTGACCACTCATTTACTTGTACAGCTCGTCCATGC | |
| 53 | CCGGAAGCGCACACTGACTTTATTTTCGCCTGCATCGGCGAGGAGCTGGGTTATGTCGGTGTGGTGCTGGCACTTTTAATG | Mutagenesis of chromosomal *ftsW* gene to *ftsW*$^{I302C}$ |
| 54 | CGCGTGTGCTTGGCTTCGCACGAGACGCAATTGTCTGCCGTATTTTTGGCGCAGGGATGGCAACCGACGCCTTTTTCGTCG | Mutagenesis of chromosomal *murJ* gene to *murJ*$^{A29C}$ |
| 55 | TTTCGCCGCTGCTTTCTTGTACAGCTCGTCCATGCC | To generate *tagrfp-t-fts*I without linker and first methionine |
| 56 | AAAGCAGCGGCGAAAA | |
| 57 | GAGGAGAAATTAACTATGGTGAGCAAAGGCGAAGAAG | Amplification of the *mneongreen* gene |

| | | |
|---|---|---|
| 58 | GGAGCCAGCGGATCCTTTATACAGTTCATCCATGCCCATC | |
| 59 | AGTTAATTTCTCCTCTTTAATGAATTCTG | Amplification of the vector with *zapA* gene for pXY677 |
| 60 | GGATCCGCTGGCTCCGCTG | |

Supplementary Table 3: Quantification of septal NAM labeling.

| **Percentage of cells with labeled septa** | **Wt (N)** | ***ftsW*$^{I302C}$** | ***Δ3, ponB*$^{S247C}$** | ***Δ3, ponB*$^{S247C}$, *ftsW*$^{I302C}$** | ***murJ*$^{A29C}$** |
|---|---|---|---|---|---|
| - MTSES | 38 ± 4% (1163) | 34 ± 4% (970) | 32 ± 1% (1128) | 29 ± 5% (1200) | 32 ± 3% (836) |
| + MTSES | 32 ± 5% (1113) | 22 ± 4% (723) | 14 ± 3% (1085) | 4 ± 1% (1346) | 2.6 ± 0.1% (1031) |
| $R_A$ (+MTSES/-MTSES) | 85 ± 6% | 64 ± 5% | 42 ± % | 13 ± % | 9 ± 2% |
| **Integrated Intensity of labeled septa** | | | | | |
| - MTSES [a] (a.u) | $1.6 \pm 0.1\times10^6$ (442) | $1.1 \pm 0.1\times10^6$ (330) | $0.6 \pm 0.1\times10^6$ (361) | $0.5 \pm 0. \times10^6$ (348) | $0.69 \pm 0.03\times10^6$ (268) |
| + MTSES[a] (a.u) | $1.5 \pm 0.1\times10^6$ (356) | $0.8 \pm 0.1\times10^6$ (160) | $0.37 \pm 0.07\times10^6$ (152) | $0.18 \pm 0.05\times10^6$ (53) | $0.37 \pm 0.02\times10^6$ (27) |
| $R_B$ (+MTSES/-MTSES) | 98± 13% | 70 ± 4% | 65 ± 12% | 41 ± 12% | 54 ± 4% |
| ***Loss of total labeling* [b]** | 16 ± 15% | 56 ± 5% | 72 ± 6% | 96 ± 2% | 95.5 ± 0.5% |

Errors are the mean of standard error of three experimental repeats

Cells were grown in M9-glucose at 25°C

N is the number of cells analyzed

[a] Average cell body intensity was subtracted from the integrated intensity of the septal area. Note the median instead of mean integrated intensity was listed in the table and used for calculations due to the skewed distribution toward high intensity values in a small population of cells. See Methods and Extended Data Fig. 3.

[b] Calculated using $(1 - R_A \cdot R_B)$ .

Supplementary Table 4: FtsW dynamics in cells with different FtsZ treadmilling speeds

| FtsZ mutation | FtsZ speed[a] (nm/s), ($N_Z$) | *P_moving*[b] (%), ($N_{all}$)[e] | $T_s$(s)[c] ($N_S$)[e] | $V_1$[d] (nm/s), ($N$)[e] | $P_1\_V_1$[d] (%) | $V_2$[d] (nm/s) | *FtsW speed*[b] (nm/s) |
|---|---|---|---|---|---|---|---|
| wt | 28.0±1.2 (182) | 61.5±1.8 (519) | 19.2±1.1 (204) | 8.0±0.3 (315) | 37.4±11.2 | 31.9±4.4 | 22.5±1.6 |
| E238A | 23.8±2.9 (37) | 61.1±2.7 (155) | 22.3±1.7 (65) | 7.9±0.4 (90) | 43.1±14.5 | 28.2±9.5 | 18.8±9.4 |
| E250A | 17.4±1.6 (41) | 61.4±2.0 (233) | 24.7±1.5 (96) | 8.4±0.3 (137) | 71.2±14.8 | 27.7±6.8 | 12.4±8.3 |
| D269A | 14.3±1.3 (32) | 50.1±2.3 (158) | 23.8±1.7 (84) | 7.8±0.4 (74) | 46.7±22.6 | 15.7±5.5 | 11.0±7.1 |
| G105S | 9.7±1.0 (35) | 47.9±2.1 (219) | 26.3±1.5 (112) | 7.6±0.5 (102) | 34.5±12.7 | 11.0±1.8 | 10.3±7.6 |
| D158A | 7.4±0.7 (36) | 39.2±2.7 (144) | 30.6±2.4 (85) | 9.0±2.9 (59) | 44.9±22.5 | 12.8±3.7 | 11.3±9.1 |

All the experiments were performed with cells growing on M9-glucose medium

[a] FtsZ treadmilling speeds are from[2]. mean ± S.E.M, where S.E.M is the standard error of mean speed. $N_z$ is the number of FtsZ kymograph segments.

[b] Percentage of time (*P_moving*) and average speed of FtsW-RFP molecules spent in directional moving state. Data expressed as mean ± error, where the error is the standard deviation from 1000 bootstrap samples pooled from three independent experiments.

[c] Average dwell time of stationary FtsW-RFP molecules. Data expressed as mean ± S.E.M.

[d] Speed ($V_1$), percentage ($P_1\_V_1$) of the slow-moving population and speed ($V_2$) of the fast-moving population of FtsW-RFP obtained from two-population free-float fitting of CDF curves bootstrapped 1000 times from three independent experiments. Errors are the standard deviations of the fitted parameters.

[e] $N_{all}$ is the number of total track segments while $N$ is the number of segments corresponding to a directionally moving molecule and $N_s$ is the number of segments corresponding to a stationary molecule.

Supplementary Table 5: FtsI dynamics in different strains

| genotype | medium | *P_moving*[a] (%), ($N_{all}$)[c] | $V_1$[b] (nm/s), ($N$)[c] | $P_1\_V_1$[b] (%) | $V_2$[b] (nm/s) | *FtsI average speed*(nm/s) |
|---|---|---|---|---|---|---|
| BW25113 | M9-glucose | 54.2±3.7 (92) | 9.8±1.1 (52) | 53.9±19.9 | 31.2±5.6 | 20.2±2.0 |
| BW25113, *ftsZ*[E250A] | M9-glucose | 58.0±4.7 (84) | 9.1±1.8 (45) | 21.9±22.6 | 19.2±3.4 | 16.5±1.7 |
| TB28, *ftsB*[E56A] | M9-glucose | 52.2±3.4 (146) | 9.2±0.7 (76) | 89.8±4.4 | 37.3±3.8 | 12.4±1.4 |

[a] Percentage of time (*P_moving*) and average speed of RFP-FtsI molecules spent in directional moving state. Data expressed as mean ± error, where the error is the standard deviation of 1000 bootstrap samples from three experiments.

[b] Speed ($V_1$), percentage ($P_1\_V_1$) of the slow-moving population and speed ($V_2$) of the fast-moving population of RFP-FtsI obtained from two-population free-float fitting of 1000 CDF curves bootstrapped from three experiments. Errors are the standard deviations of the fitted parameters.

[c] $N_{\mathrm{all}}$ is the number of total track segments while $N$ is the number of segments corresponding to a directionally moving molecule.

Supplementary Table 6: FtsW dynamics under different sPG synthesis conditions

| genotype | Drug or medium | *P_moving*[b] (%), ($N_{all}$)[f] | $T_s$[c](s), ($N_s$)[f] | $V_1$[c] (nm/s), (*N*)[f] | $P_1$_$V_1$[c] (%) | $V_2$[c] (nm/s) | *FtsZ speed*[e] (nm/s), ($N_z$) | *FtsW average speed*(nm/s) |
|---|---|---|---|---|---|---|---|---|
| BW25113 | EZRDM | 63.6±0.8 (1445) | 13.6±0.5 (535) | 11.1±0.24 (910) | 80.7±1.8 | 51.9±2.3 | 49.8±2.9 (182) | 18.6±0.8 |
| BW25113 | M9-glucose | 61.5±1.8 (519) | 19.2±1.1(204) | 8.0±0.3 (315) | 37.4±11.2 | 31.9±4.4 | 28.0±1.2 (182) | 22.5±1.6 |
| BW25113 | Fosfomycin, M9-glucose | 20.5±1.3 (643) | 15.2±0.8(404) | 9.6±0.1 (239) | 7.9±9.3 | 34.9±2.9 | 39.7±1.9 (192) | 31.4±1.6 |
| BW25113 | Aztreonam, M9-glucose | 11.1±3.4 (347) | 24.6±1.4(245) | 9.2±0.1 (102) | 19.8±7.7 | 34.6±2.0 | 39.2±2.1 (197) | 29.9±2.0 |
| BW25113, *ftsW*$^{I302C}$ | M9-glucose | 52.4±1.8 (361) | 18.2±1.0 (169) | 10.7±0.9 (192) | 55.4±7.5 | 33.8±2.7 | 38.6±3.2 (85) | 20.7±1.8 |
| BW25113, *ftsW*$^{I302C}$ | MTSES, M9-glucose | 38.0±1.2 (757) | 20.1±0.7 (418) | 10.4±2.5 (339) | 43.4±9.1 | 30.0±2.4 | 35.1±2.0 (85) | 22.5±1.3 |
| *MG1655,Δ3 ponB*$^{S247C}$ *ftsW*$^{I302C}$ | M9-glucose | 46.7±2.1 (499) | 11.9±0.9(223) | 9.2±0.1 (276) | 16.5±6.2 | 32.1±1.5 | 42.3±2.1 (166) | 26.5±1.5 |
| *MG1655,Δ3 ponB*$^{S247C}$ *ftsW*$^{I302C}$ | MTSES, M9-glucose | 19.6±1.8 (363) | 18.9±1.9(206) | 9.1±0.2 (157) | 1.2±2.7 | 32.1±1.5 | 38.1±2.8 (98) | 33.9±2.3 |
| TB28 | M9-glucose | 44.8±1.7 (477) | 18.2±0.8(258) | 9.4±0.3 (219) | 63.6±7.6 | 37.8±6.1 | ND[a] | 18.8±1.6 |
| MG1655, $P_{BAD}$::*ftsN* | M9-glucose [d] | 31.4±1.0 (1094) | 16.3±0.6(696) | 10.5±0.8 (398) | 41.7±14.7 | 30.1±3.9 | ND[a] | 22.1±1.3 |
| TB28, *ftsB*$^{E56A}$ | M9-glucose | 36.3±1.6 (499) | 20.3±0.8(323) | 9.3±0.8 (176) | 93.3±3.7 | 46.9±8.4 | 50.2±2.4 (180) | 11.7±1.1 |
| TB28, *ΔftsN*, *ftsB*$^{E56A}$ | M9-glucose | 36.0±1.7 (339) | 23.4±1.3 (195) | 7.9±0.6 (144) | 76.2±4.4 | 56.2±2.7 | ND[a] | 18.2±2.3 |
| TB28, *ftsW*$^{E289G}$ | M9-glucose | 49.8±1.7 (383) | 18.2±1.0 (107) | 8.9±0.9 (176) | 79.1±5.6 | 35.4±3.7 | ND[a] | 13.9±1.1 |

| | | | | | | | | |
|---|---|---|---|---|---|---|---|---|
| TB28, *ftsI*[R167S] | M9-acetate | 35.6±1.3 (756) | 19.4±0.6(496) | 6.2±0.5 (260) | 55.6±4.8 | 28.7±2.7 | 31.6±2.0 (84) | 16.8±1.5 |
| TB28, *ftsI*[R167S] | M9-glucose | 60.6±1.3 (491) | 17.6±0.7(236) | 6.4±0.2 (255) | 90.7±2.1 | 37.2±1.0 | 31.6±2.0 (92) | 8.8±0.6 |
| TB28, *ftsI*[R167S] | EZRDM | 60.1±2.2 (173) | 12.0±0.9(81) | 11.3±0.7 (92) | 87.5±5.5 | 40.0±1.5 | 37.2±2.8 (83) | 14.5±1.4 |
| TB28, *ftsI*[R167S] | EZRDM+UppS | 55.7±1.1 (954) | 11.1±0.4(441) | 13.2±0.3 (513) | 93.9±2.2 | 40.0±1.4 | ND[a] | 14.5±0.4 |
| TB28, *ftsI*[R167S] | 1XPBS (4% PFA fixed) | 2.3±0.4 (182) | NA[a] | NA[a] | NA[a] | NA[a] | NA[a] | 33.6±3.3 |

[a] N.A. not applicable. N.D. not measured

[b] Percentage of time (*P_moving*) and average speed of FtsW-RFP molecules spent in directional moving state. Data expressed as mean ± error, where the error is the standard deviation of 1000 bootstrap samples from three experiments.

[c] Speed ($V_1$), percentage ($P_1\_V_1$) of the slow-moving population and speed ($V_2$) of the fast-moving population of FtsW-RFP obtained from two-population free-float fitting of 1000 CDF curves bootstrapped from three experiments. Errors are the standard deviations of the fitted parameters.

[d] Medium without arabinose (FtsN-depleted).

[e] FtsZ's treadmilling speed, mean ± S.E.M, where S.E.M is the standard error of mean speed. $N_z$ is the number of FtsZ kymograph segments.

[f] $N_{all}$ is the number of total track segments while $N$ is the number of segments corresponding to a directionally moving molecule and $N_s$ is the number of segments corresponding to a stationary molecule.

Supplementary Table 7. Superfission variants of FtsB, FtsI or FtsW cause a short-cell phenotype, and can rescue Δ*ftsN* cells.

| Row[a] | Strain | Genotype | *Td*[b] | *N*[c] | Average length | | Average width | | Average volume | |
|---|---|---|---|---|---|---|---|---|---|---|
| | | | | | µm (SD) | %WT | µm (SD) | %WT | µm (SD) | % WT |
| LB | | | | | | | | | | |
| 1 | TB28 | wt | 41 | 303 | 4.42 (1.07) | 100 | 0.95 (0.06) | 100 | 2.91 (0.76) | 100 |
| 2 | BL167 | *ftsB*$^{E56A}$ | 45 | 324 | 3.42 (0.82) | 77 | 1.05 (0.07) | 110 | 2.65 (0.71) | 91 |
| 3 | BL173 | *ftsB*$^{E56A}$ Δ*ftsN* | 45 | 235 | 9.87 (4.56) | 235 | 0.92 (0.05) | 97 | 6.38 (3.19) | 219 |
| 4 | PM17 | *ftsW*$^{E289G}$ | 43 | 331 | 3.40 (0.74) | 81 | 1.00 (0.07) | 105 | 2.40 (0.65) | 82 |
| 5 | PM18 | *ftsW*$^{E289G}$ Δ*ftsN* | 43 | 343 | 6.28 (1.79) | 142 | 0.96 (0.05) | 101 | 4.27 (1.32) | 147 |
| 6 | PM6 | *ftsI*$^{R167S}$ | 43 | 283 | 3.53 (0.80) | 80 | 0.98 (0.07) | 103 | 2.40 (0.63) | 82 |
| 7 | PM8 | *ftsI*$^{R167S}$ Δ*ftsN* | N.A[d]. Cells not viable | | | | | | | |
| M9-glucose | | | | | | | | | | |
| 8 | TB28 | wt | 99 | 333 | 2.62 (0.57) | 100 | 1.01 (0.08) | 100 | 1.82 (0.46) | 100 |
| 9 | BL167 | *ftsB*$^{E56A}$ | 101 | 363 | 2.10 (0.44) | 80 | 1.08 (0.08) | 107 | 1.60 (0.39) | 88 |
| 10 | BL173 | *ftsB*$^{E56A}$ Δ*ftsN* | 101 | 288 | 5.45 (2.24) | 208 | 0.94 (0.06) | 93 | 3.56 (1.46) | 196 |
| 11 | PM17 | *ftsW*$^{E289G}$ | 98 | 307 | 2.41 (0.58) | 92 | 1.06 (0.07) | 105 | 1.80 (0.47) | 99 |
| 12 | PM18 | *ftsW*$^{E289G}$ Δ*ftsN* | 93 | 349 | 4.56 (1.06) | 174 | 0.97 (0.08) | 96 | 3.15 (0.88) | 173 |
| 13 | PM6 | *ftsI*$^{R167S}$ | 100 | 336 | 2.20 (0.44) | 84 | 1.06 (0.09) | 105 | 1.60 (0.39) | 88 |
| 14 | PM8 | *ftsI*$^{R167S}$ Δ*ftsN* | N.A[d], cells are viable but grow poorly and are very filamentous | | | | | | | |

[a] Cells were grown at 30°C. Overnight cultures were diluted to $OD_{600}$ = 0.02 in LB and grown to $OD_{600}$ = 0.6-0.7 (rows 1-6), or to $OD_{600}$ = 0.05 in M9-glucose and grown to $OD_{600}$ = 0.5-0.6 (rows 8-13). Cells were then chemically fixed, imaged with phase-contrast optics, and measured using MicrobeJ software[28].

[b] Mass doubling time in minutes.

[c] Number of cells measured.

[d] N.A. not available

Supplementary Table 8. Growth and constriction rates of TB28, *ftsI*$^{R167S}$ cells.

| medium | *Doubling Time*[a] (min) | *Constriction rate* [c] (nm/min), (*N*)[d] |
|---|---|---|
| M9-acetate | 806±20[b] | 4.9±1.1[e] (5) |
| M9-glucose | 202±9 | 17.2±1.8 (15) |
| M9-EZRDM | 57.2±3.8 | 37.9±2.0 (16) |
| M9-EZRDM+UppS | NA | 39.7±2.3 (20) |

[a] Mass doubling time in minutes at 25°C in liquid culture.

[b] mean ± S.E.M from 2-3 independent measurements.

[c] Constriction rates on gel-pad at ~ 25°C

[d] Number of cells measured.

[e] mean ± S.E.M.

Supplementary Table 9. Mean Error of single- and double-population fitting of CDF curves.

| genotype | Drug or medium | *Error_1*[a] | *Error_2*[b] | genotype | Drug or medium | *Error_1*[a] | *Error_2*[b] |
|---|---|---|---|---|---|---|---|
| FtsW-RFP | | | | | | | |
| BW25113 | EZRDM | 0.012 | 0.0031 | BW25113, ftsZE238A | M9-glucose | 0.015 | 0.0089 |
| BW25113 | M9-glucose | 0.014 | 0.0037 | BW25113, ftsZE250A | M9-glucose | 0.010 | 0.0052 |
| BW25113 | Fosfomycin, M9-glucose | 0.0060 | 0.0051 | BW25113, ftsZD269A | M9-glucose | 0.013 | 0.0088 |
| BW25113 | Aztreonam, M9-glucose | 0.0081 | 0.0049 | BW25113, ftsZG105S | M9-glucose | 0.010 | 0.0066 |
| BW25113, *ftsW*$^{I302C}$ | M9-glucose | 0.0122 | 0.0040 | BW25113, ftsZD158A | M9-glucose | 0.011 | 0.0075 |
| BW25113, *ftsW*$^{I302C}$ | MTSES, M9-glucose | 0.0060 | 0.0033 | MG1655, PBAD::ftsN | M9-glucose | 0.0046 | 0.0039 |
| *MG1655,Δ3 ponB*$^{S247C}$ *ftsW*$^{I302C}$ | M9-glucose | 0.0070 | 0.0037 | TB28, ftsBE56A | M9-glucose | 0.0079 | 0.0048 |
| *MG1655,Δ3 ponB*$^{S247C}$ *ftsW*$^{I302C}$ | MTSES, M9-glucose | 0.0062 | 0.0046 | TB28, ΔftsN, ftsBE56A | M9-glucose | 0.0160 | 0.0061 |
| TB28 | M9-glucose | 0.0125 | 0.0047 | TB28, *ftsW*$^{E289G}$ | M9-glucose | 0.0098 | 0.0048 |
| TB28, *ftsI*$^{R167S}$ | M9-glucose | 0.0105 | 0.0032 | TB28, *ftsI*$^{R167S}$ | EZRDM | 0.0103 | 0.0051 |
| TB28, *ftsI*$^{R167S}$ | M9-acetate | 0.0130 | 0.0048 | TB28, *ftsI*$^{R167S}$ | EZRDM +UppS | 0.0063 | 0.0051 |
| RFP-FtsI | | | | | | | |
| BW25113 | M9-glucose | 0.013 | 0.0060 | BW25113, ftsZE250A | M9-glucose | 0.010 | 0.0058 |
| TB28, ftsBE56A | M9-glucose | 0.011 | 0.0057 | | | | |

[a] mean CDF fitting error of single population log-normal distribution. The mean error is calculated as:

$$\langle Error \rangle = < \frac{1}{N}\sum_{i=1}^{N}\sqrt{R_i^2} >$$

Where $R_i$ is the residual of data point i of the CDF curve.

[b] mean CDF fitting error of double-population log-normal distribution. The mean error is calculated as in[a].

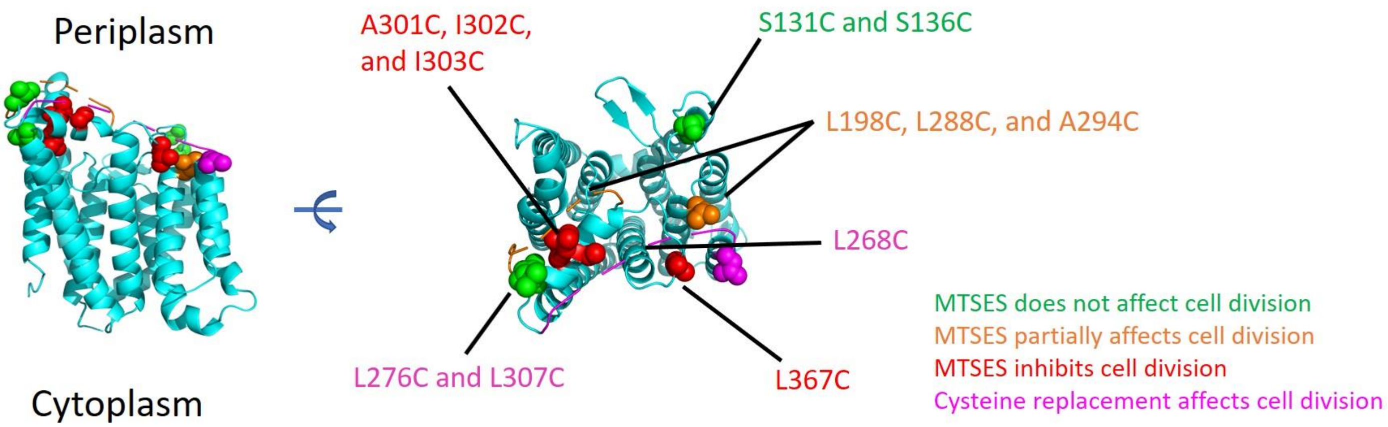


**Extended Data Figure 1**. **Homology structure of FtsW based on RodA structure (PDB: 6BAR)** [29]**using Phyre 2.0**[30]**.**

Mutated residues are labeled and color-coded according to their sensitivity to MSTES. Green: MTSES does not affect cell growth or morphology. Orange: MTSES significantly slows down cell division and leads to elongated cells. Red: MTSES blocks cell division completely and leads to long, chaining cells. Magenta: Cysteine mutation causes other cell division defects such as abnormal septa and cell poles even in the absence of MTSES.

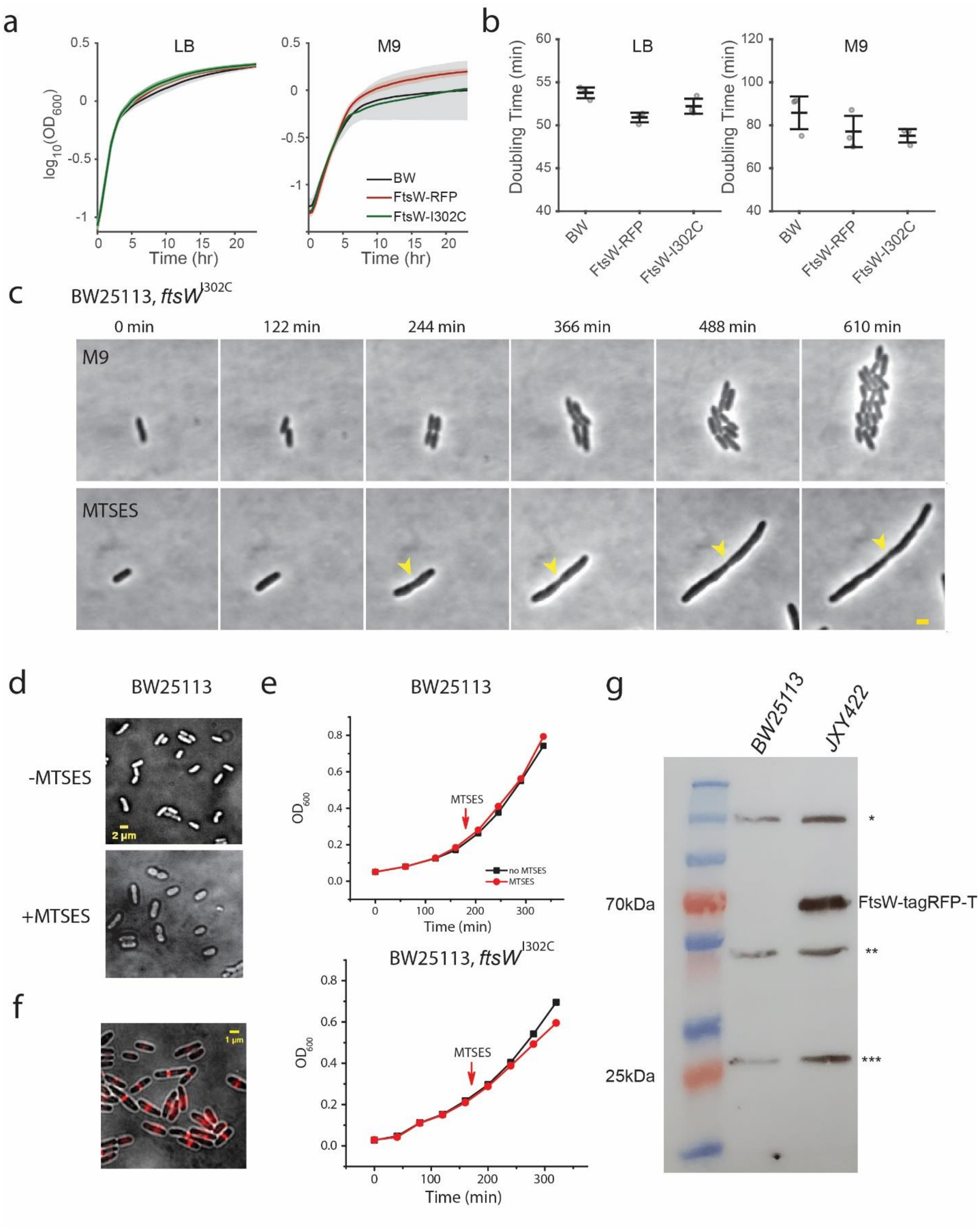


**Extended Data Figure 2**. **Characterizations of FtsW, FtsW$^{I302C}$ and FtsW-TagRFP-t fusion proteins used in the study.**

**a.** Growth curves of BW25113 (*wt*), JXY422 (*ftsW-TagRFP-t*), and JXY559 (*ftsW*$^{I302C}$) cells in LB or M9-glucose medium **b**. Doubling times of the three strains in LB and M9 calculated from growth curves in **a**. **c**. Time-lapse images of JXY559 (*ftsw*$^{I302C}$) cells in M9 medium without (top panel) and with (bottom panel) MTSES. MTSES was added to the cell in the bottom panel 30 min prior to the first image (Supplementary Movie 1 and 2). Arrowhead marks a cell constriction that initiated early on but failed to complete. Scale bar:2 µm. **d**. Snapshots of log phase BW25113 (*wt*) cells grown in M9 medium without (top panel) or with (bottom) MTSES. Scale bar:2 µm. **e**. Growth curves of BW25113 (*wt*) and JXY559 (*ftsW*$^{I302C}$) cells in M9 medium upon addition of MTSES in early log-phase. **f**. JXY422 (*ftsW-TagRFP-t*) cells showing the expected midcell localization of FtsW-TagRFP-t. Scale bar: 1µm. **g**. Immunoblot of FtsW-TagRFP-t using anti-TagRFP antibody. As expected, the FtsW-TagRFP-t band (~70kDa) is present in JXY422 but not in BW25113. Three non-specific bands (*, **, ***) are evident in both samples. Band*** migrates close to what would be expected of free TagRFP-t. If the modestly increased intensity of band*** in the JXY422 sample were due to degradation of the full length FtsW-TagRFP-t, it would represent only a minor fraction (< 4%) of total FtsW-TagRFP-t. Cells were grown at room temperature (c, f, g), 30$^{o}$C (a, b), or 37$^{o}$C (e). MTSES was used at 0.1 mM in c and d; 1 mM in e.

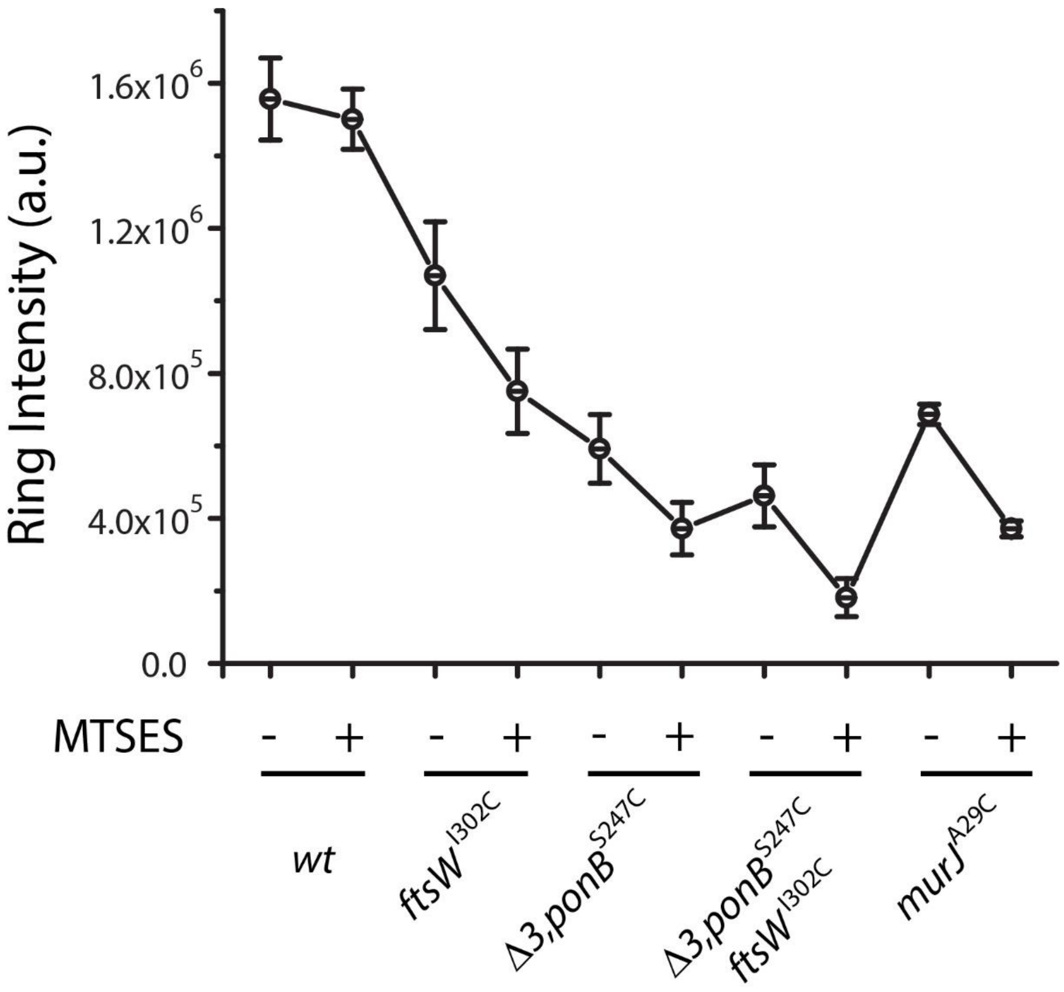


**Extended Data Figure. 3**. **Integrated intensity of septal NAM labeling of different strains in the absence or presence of MTSES**. Error bars represent S.E.M of three independent experiments with the same imaging condition.

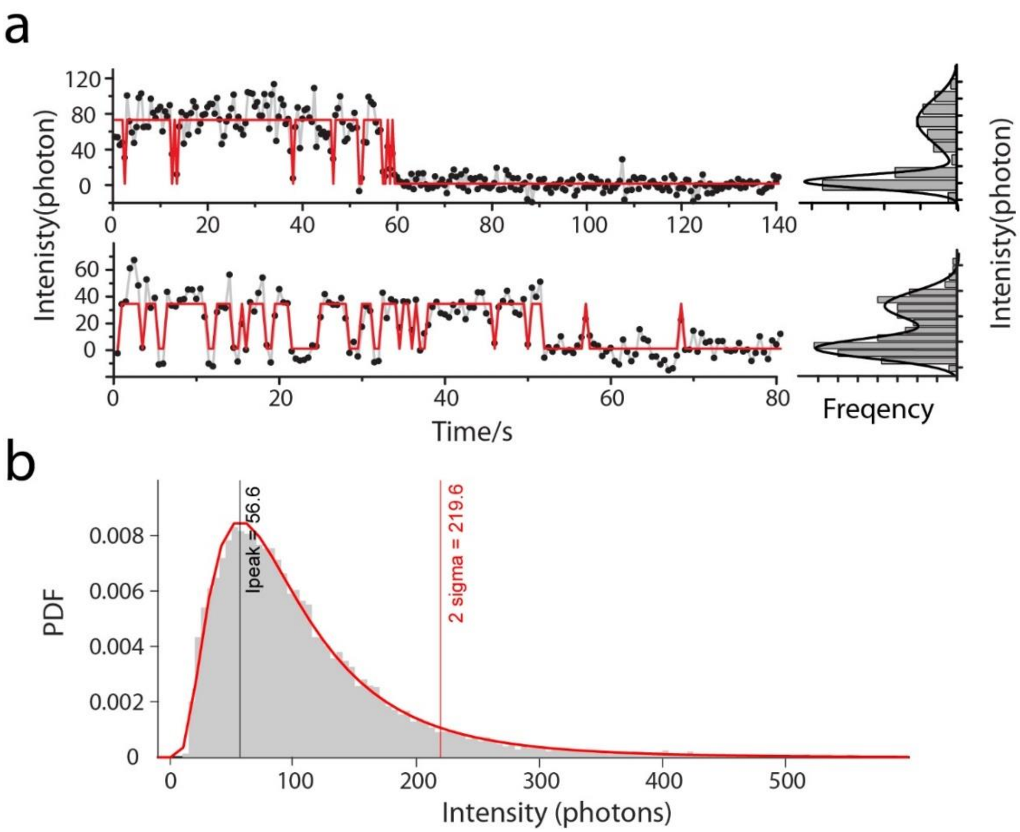


**Extended Data Figure.** 4. **Example of single-particle intensity traces and allover intensity distribution of FtsW-RFP in BW25113 strain in M9-glucose medium.**

a. Intensity traces show single step photobleach and blinking behavior, which suggest they are single FtsW-RFP molecules. Intensity was subtracted by the average background and integrated over a 5-by-5 pixel region, and back-calculated to photon numbers using experimental camera setting (EmGain = 300, preAmp = 9.93, Quantum yield ~ 0.96). Intensity histograms are shown on the right.

b. Probability density function (PDF) of all fluorescent spots in BW25113 strain expressing FtsW-RFP. Most of the spots are from single FtsW-RFP (~56 photons). To minimize the possibility of tracking multiple FtsW-RFP molecules at a time, we removed spots with intensity > 2 sigma (~4 molecules) prior to further analysis.

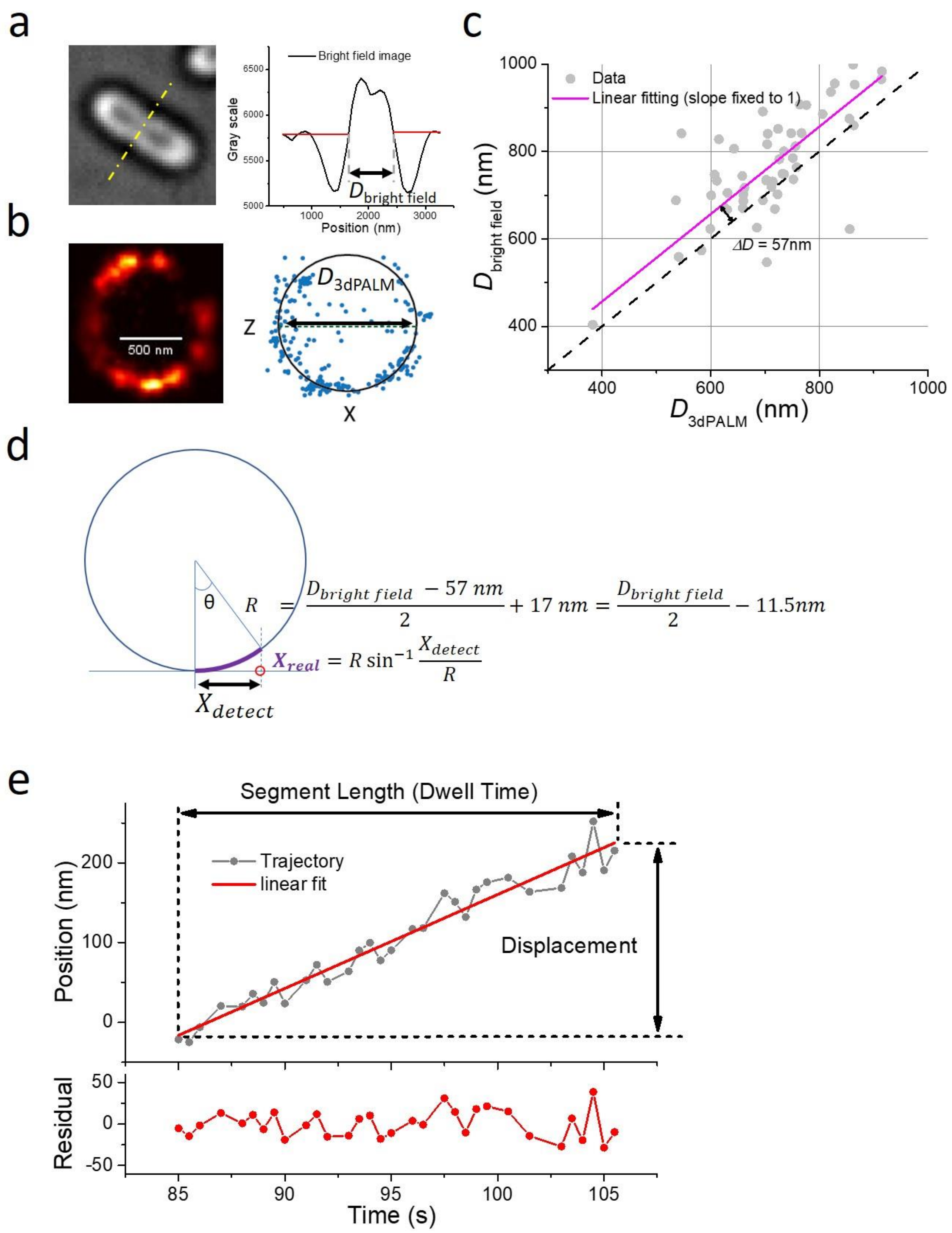


**Extended Data Figure 5**. **Custom-developed cell envelope unwrapping method to retrieve true coordinates of single molecules along the circumference of the curved cell surface**.

**a**. The apparent cell diameter ($D_{bright\ field}$, right panel) is measured using the intersections of the background signal (red line) with the cell profile (black curve) resulted from a line scan (yellow dashed line) of the cell's bright-field image (left panel). **b**. For each cell (BW25113/pJL005), the corresponding diameter of FtsZ ring ($D_{3dPALM}$) is measured using three-dimensional (3D) superresolution image of FtsZ-mEos3.2 (left) fit with a circle (right). **c**. The apparent cell diameter $D_{bright\ field}$ is plotted against $D_{3dPALM}$ from the same cell and fit with a line with a slope of 1 and intersection at 57nm (magenta line). **d**. The true cell radius (*R*, from cell center to the inner membrane) is calculated from $D_{bright\ field}$ subtracting the intersection and adding the distance between FtsZ ring and inner membrane (~ 17nm[31]). The 'true' coordinate $X_{\mathrm{real}}$ (purple arc) of a molecule along the circumference of the cell inner membrane is calculated using the true cell radius R and the detected *X* coordinate $X_{\mathrm{detect}}$ (bottom equation). **e**. One segment of a single FtsW-RFP trajectory along the circumference fits to a line. The displacement (*L*) and the segment length (*T*) introduced in the Methods are labeled in the top panel. The noise level *S* is defined as the standard deviation of the residuals ($s_{\mathrm{i}}$, bottom panel, Methods).

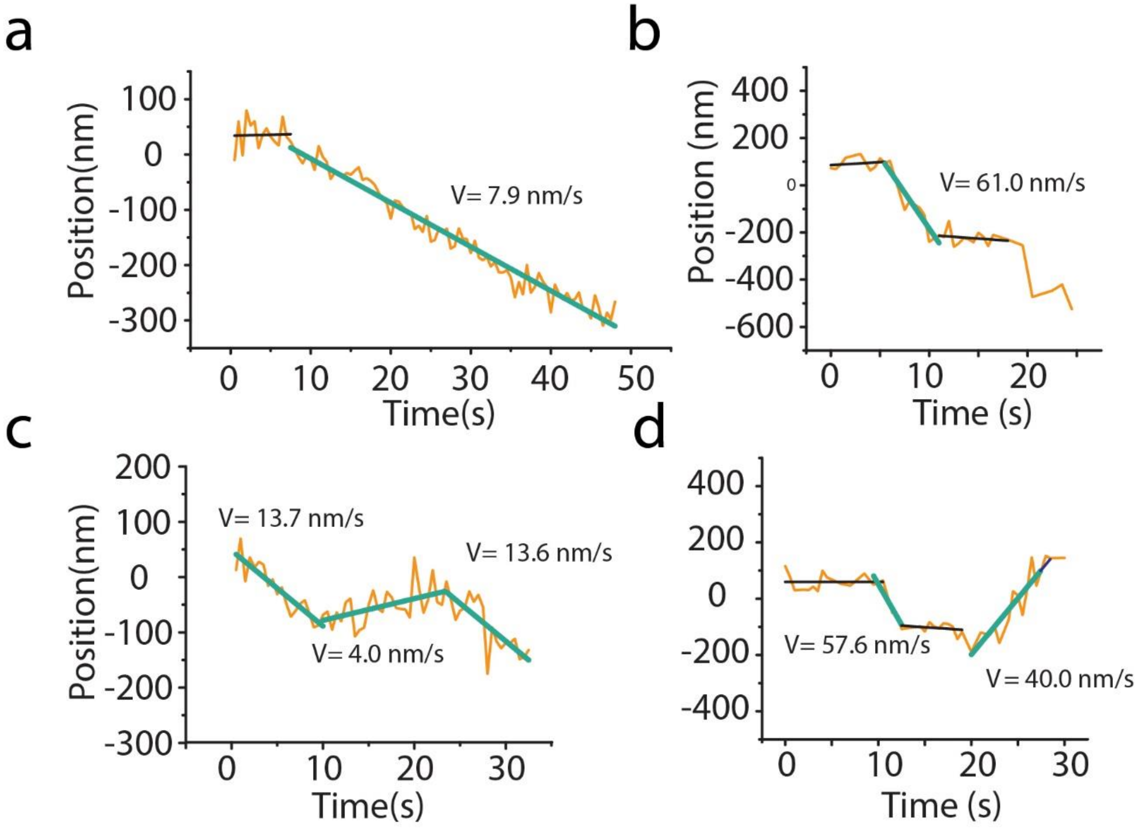


**Extended Data Fig. 6**. **Representative circumference trajectories (yellow) of FtsW-RFP molecules at septa are segmented and fit with line to extract directional moving speeds**.

Thick green lines indicate segments of directional movement. Corresponding speeds, obtained by linear fitting, are shown. Back lines indicate stationary segments. Only segments representing more than five data points (2.5 seconds) were used for fitting. The stationary segments are indicated by the gray lines.

Supplementary Movies 6 to 9 correspond to trajectories a to d.

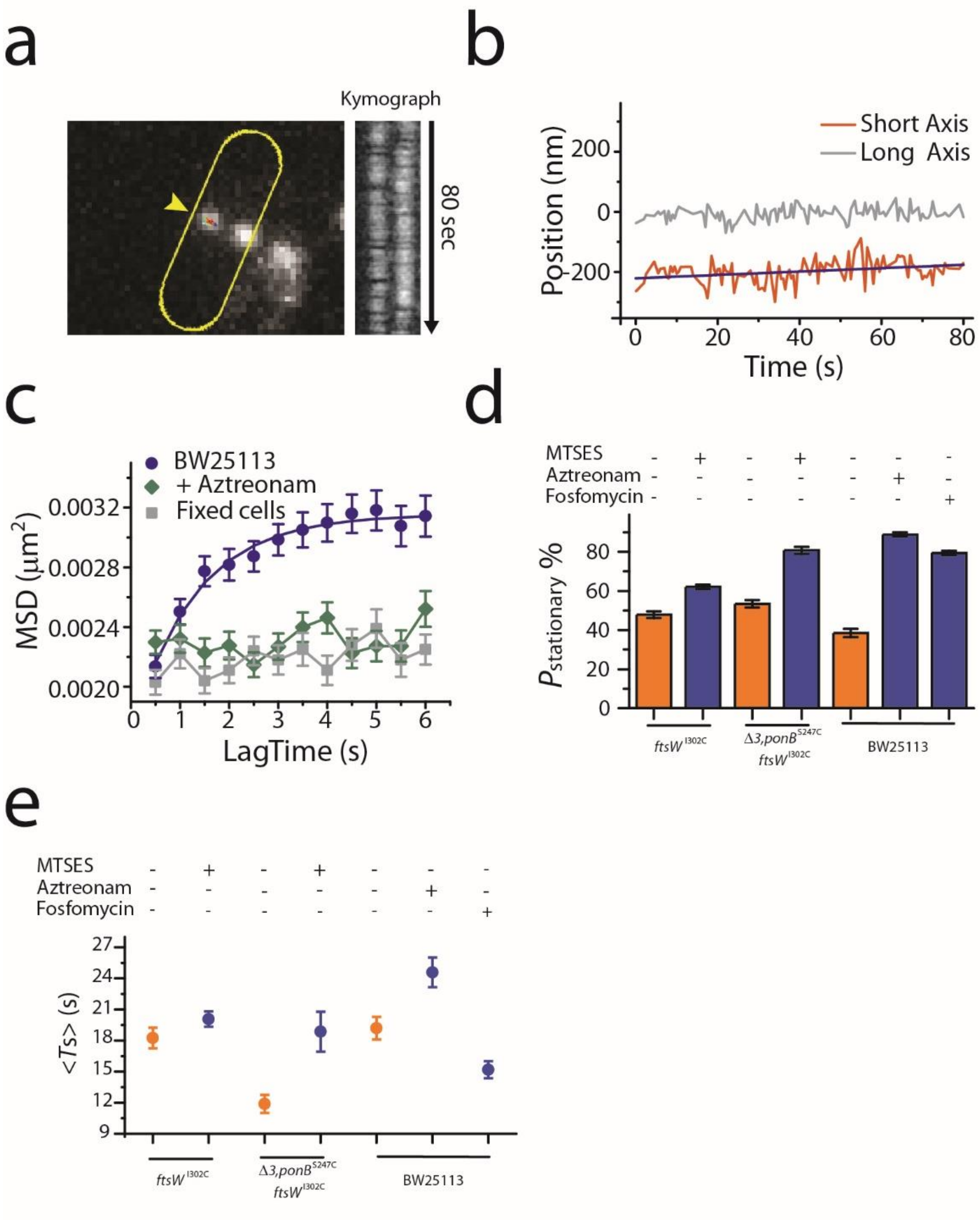


**Extended Data Figure. 7. Characterization of FtsW molecules in stationary state or confined diffusion.**

**a**. Example of an FtsW molecule (arrowhead) remaining within a small region of the septum. As indicated by the kymograph, this molecule showed little observable directional movement for at least 80 seconds. **b**. Unwrapped and decomposed 1d-trajectory of the FtsW molecule along the long (gray) and short (orange) axis. **c**. One dimensional MSD curves (short axis) of 204 ‘stationary’ molecules in live BW25113 [*wt*] cells growing in M9-glucose medium without (blue) or with 1µg/mL Aztreonam (dark green), and in cells grown without Aztreonam upon fixation with para-formaldehyde and suspension in PBS (gray). The BW25113 curve (blue) is fit by Kusumi equation[32] with D = 0.0007 $\mu m^2/s$, L = 95.5 nm. Note that Aztreonam treatment causes confinement of ‘stationary’ FtsW-RFP molecules to smaller areas, similar to what is observed in fixed cells. **d**. Percentage of FtsW-RFP molecules in stationary state in the presence (blue bars) and absence (orange bars) of drug to inhibit sPG synthesis (MTSES for $FtsW^{I302C}$ and $PBP1B^{S247C}$, Aztreonam for PBP3, and Fosfomycin for MurA). **e**. Mean dwell time of FtsW-RFP molecules staying in stationary state in the presence (blue dots) and absence (orange dots) of drug to inhibit sPG synthesis, as in panel d.

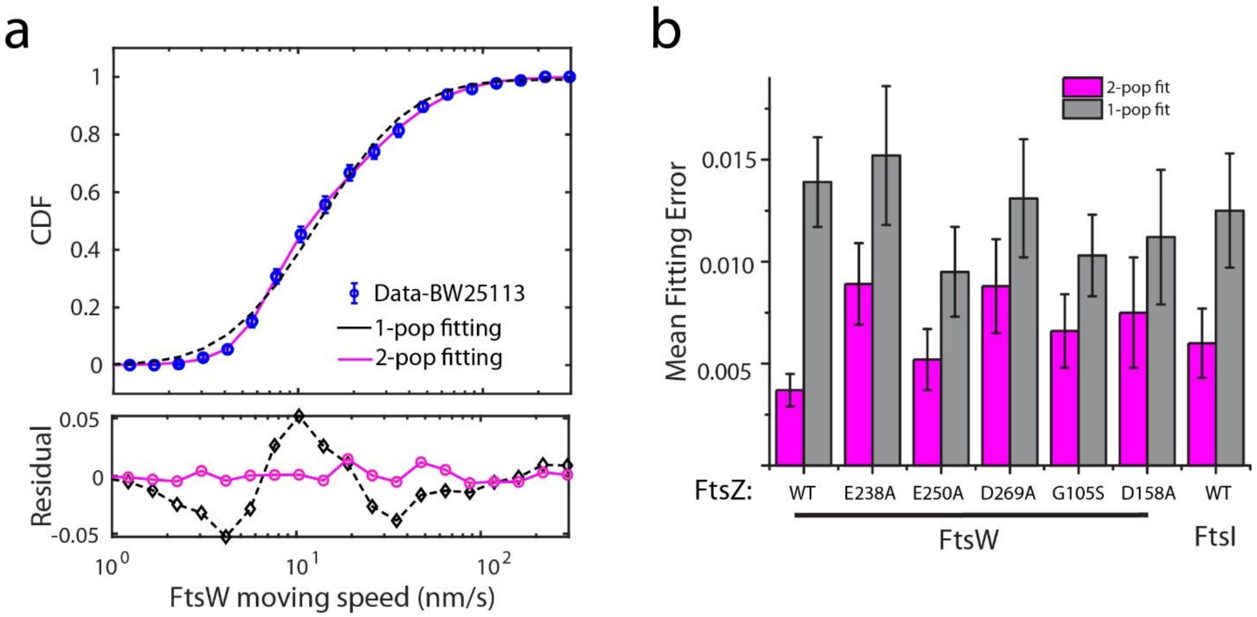


**Extended Data Fig. 8**: **Single- or double-population fitting of the cumulative probability density of directional moving speed distribution**.

**a**. CDF curve of the directional moving speed of FtsW-RFP molecules in WT BW25113 cells (blue circles) was best fit by a double (solid magenta curve) instead of a single (dash gray curve) population (empirically using log-normal distribution to describe the long tail), as indicated by the residuals below. **b.** Standard error of the mean of the fitting residuals from 1000 times bootstrapping (bottom panel in a) indicates that most CDF curves were best fit with two populations (magenta) in both BW25113 and FtsZ GTPase mutant cells.

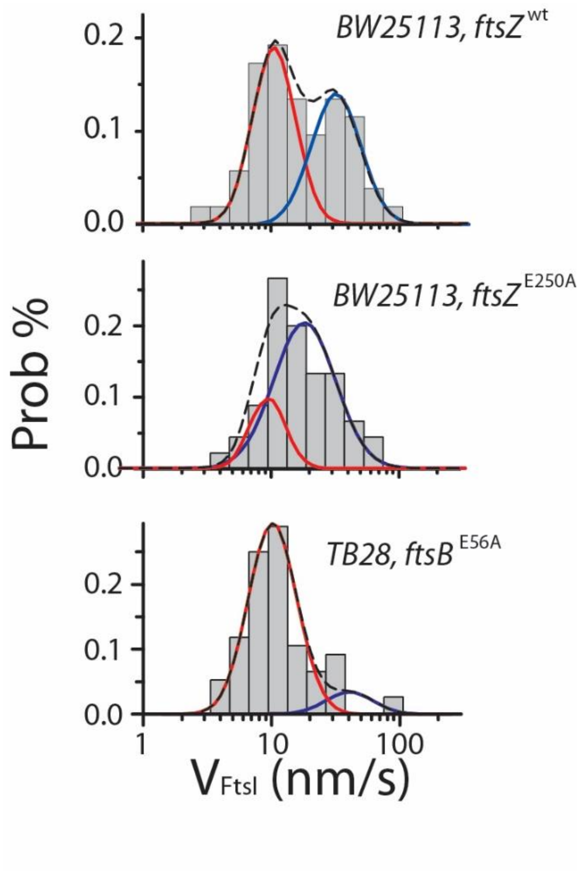


**Extended Data Fig. 9**: **Speed distributions of processive moving RFP-FtsI molecules.**

Speed distribution (bars) of all RFP-FtsI molecules in *ftsZ* wt, GTPase mutant (E250A), and *ftsB* super-fission mutant (E56A) strains. The CDF fit curves of fast- (blue solid) and slow- (red solid) moving population are overlaid with the bar graph. The black dash curves indicate the overall fitting (The fitting results are listed in Supplementary Table 5).

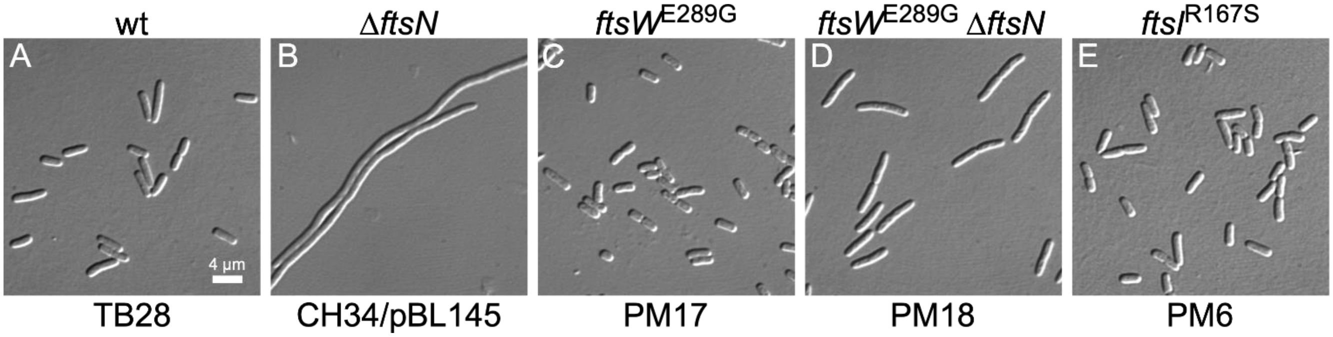


**Extended Data Fig. 10. The E289G substitution in FtsW allows for growth and division in the absence of FtsN.**

Differential interference contrast (DIC) cell images of strains TB28 (wt) (**A**), CH34/pBL145 (Δ*ftsN* / *cI*$^{ts}$ $P_{\lambda R}$::*ftsN*$^{1\text{-}90}$) (**B**), PM17 (*ftsW*$^{E289G}$) (**C**), PM18 (*ftsW*$^{E289G}$ Δ*ftsN*) (**D**), and PM6 (*ftsI*$^{R167S}$) (**E**). Cells were grown overnight in LB medium at 30$^o$C (A, C-E) or 37$^o$C (B), diluted to $OD_{600}$ = 0.02 in LB, and grown to $OD_{600}$ = 0.6-0.7 at 30$^o$C (causing depletion of FtsN in the cells shown in B). Scale bar: 4 µm.

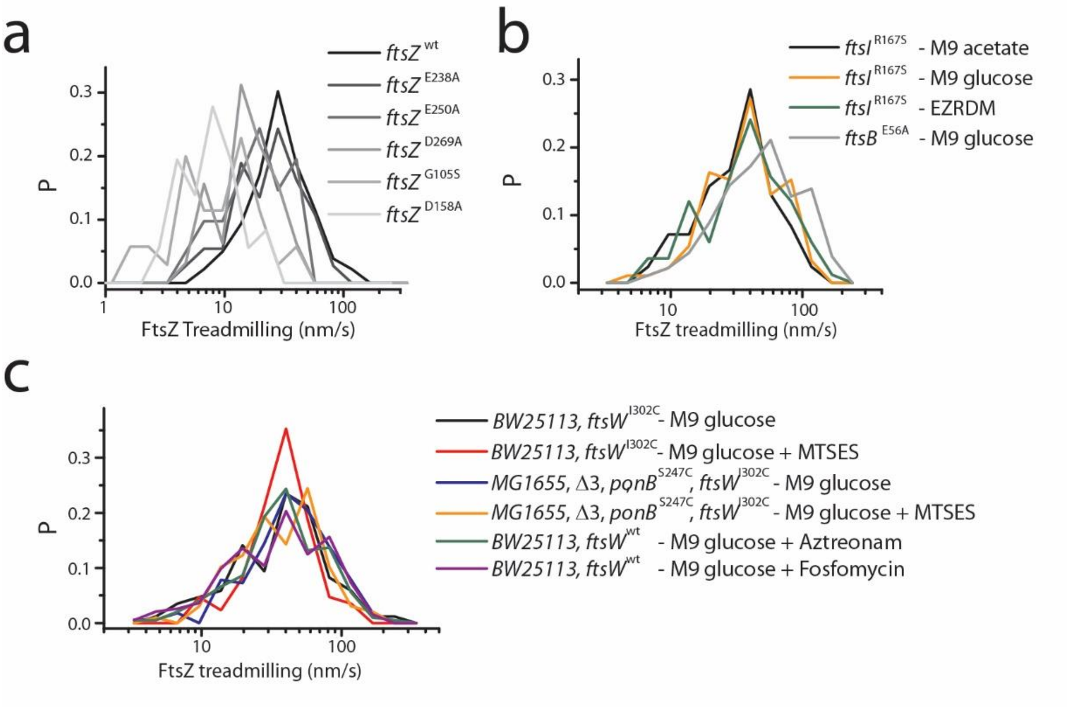


**Extended Data Fig. 11. The speed distributions of FtsZ treadmilling.**

**a.** FtsZ treadmilling speed distribution in different FtsZ GTPase mutant strains (Data from[2]). **b.** FtsZ treadmilling speed distributions in super-fission mutant strains and different growth media. **c.** FtsZ treadmilling speed distributions under different drug treatment conditions. The average speeds under all conditions are given in Supplementary Tables 4 and 6.

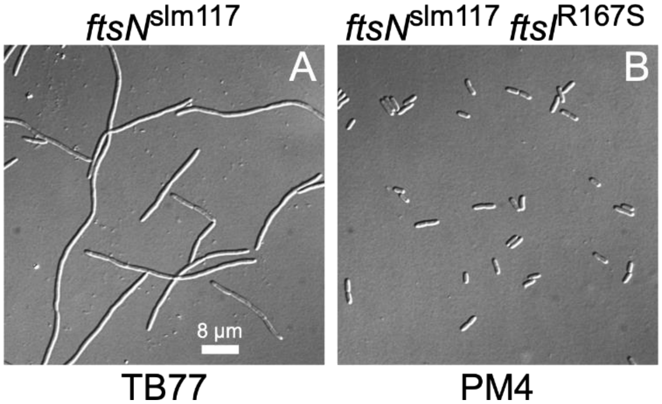


**Extended Data Fig. 12. Stimulation of constriction in cells with diminished FtsN function by the *ftsI*$^{R167S}$ superfission allele.**

DIC cell images of strains TB77 (*ftsN*$^{slm117}$) (**A**), and PM4 (*ftsN*$^{slm117}$ *ftsI*$^{R167S}$) (**B**). Overnight cultures were diluted to $OD_{600}$ = 0.05 in M9-glucose medium and grown to $OD_{600}$ = 0.5-0.6 at 30$^{o}$C. Note that the *ftsN*$^{slm117}$ allele corresponds to an EZTnKan-2 transposon insertion in codon 119 of *ftsN*, leading to a pronounced, but non-lethal, cell constriction defect[13]. Also note that the *ftsI*$^{R167S}$ allele in strain PM4 largely overcomes this defect. Scale bar: 8 µm.

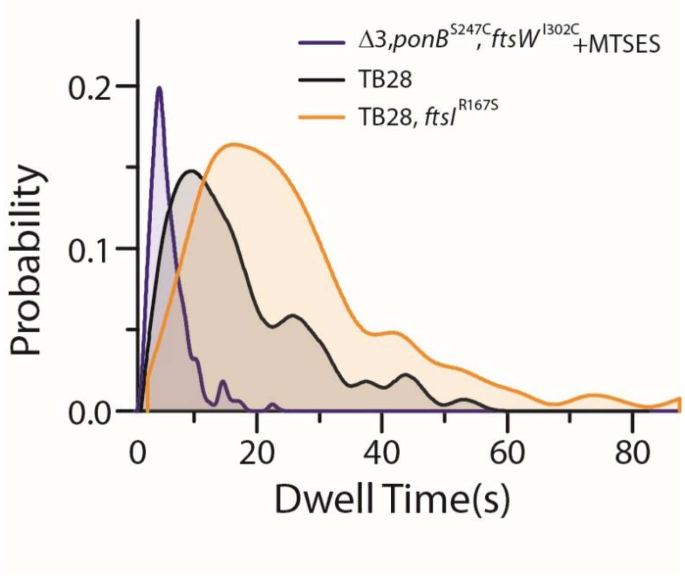


**Extended Data Fig. 13. Dwell time distribution of processive moving FtsW-RFP molecules at the division septum.**

Black curve: Dwell time distribution of processive moving FtsW-RFP molecules in wildtype TB28 cells. Orange curve: Moving FtsW-RFP molecules display longer dwell times in cells of the FtsI superfission mutant strain PM6 (TB28, $ftsI^{R167S}$). Blue curve: Inhibition of sPG synthesis (all PGTases) results in short dwell times. Note that the imaging conditions for each curve were identical, indicating that differences in these distributions were not due to different levels of photobleaching. The differences are rather more reflective of FtsW-RFP molecules spending more (orange) or less (blue) time than usual on the sPG-track (synthesizing sPG), rather than on the Z-track (tracking FtsZ filaments).

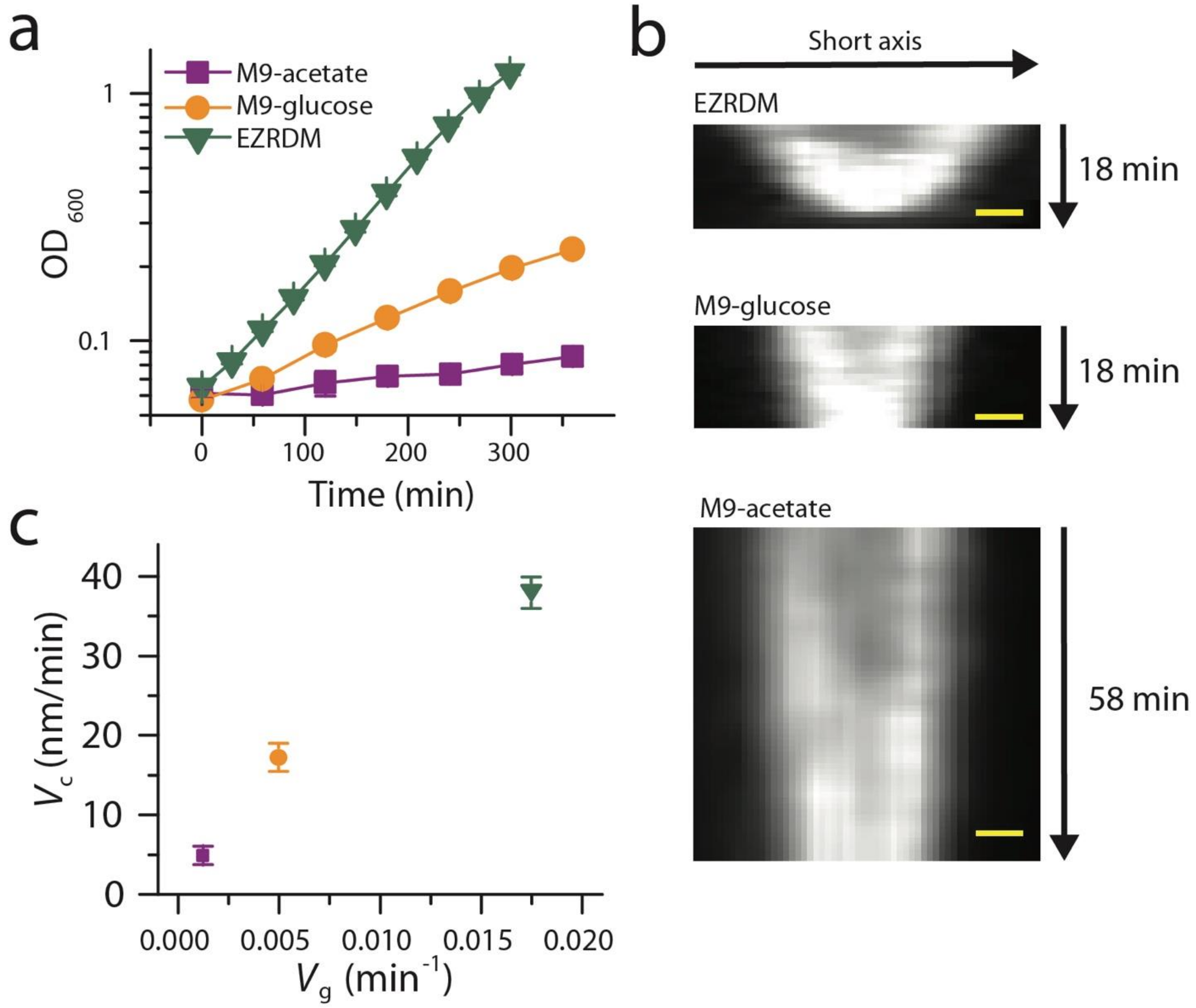


**Extended Data Fig. 14. Cell growth and constriction rate of FtsI superfission strain PM6 (TB28, *ftsI*$^{R167S}$).**

a. Growth curves of PM6 in M9-acetate (magenta), M9-glucose (orange), and EZRDM (olive).

b. Representative kymographs of the septum closure progress in PM6 cells growing in the indicated media, as probed by mNeonGreen-ZapA fluorescence. Scale bars: 200nm

c. Relationship between the cell growth rate ($V_g$, estimated from **a**) and the constriction rate ($V_c$, estimated from kymographs as in **b**). The rates are listed in Supplementary Table 8.

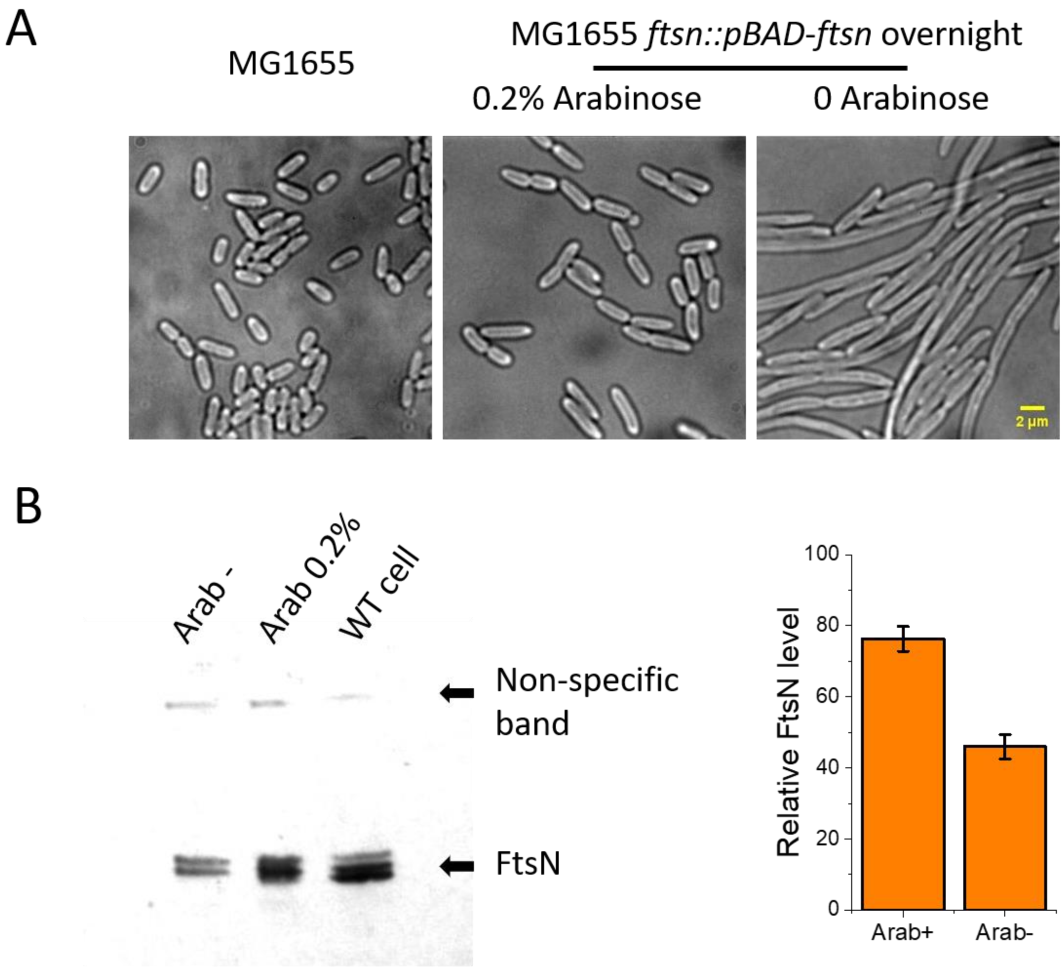


**Extended Data Fig. 15. Characterization of the FtsN depletion strain (EC1908).**

**a**. Bright field cell images of *wt* strain MG1655 strain, and of the FtsN-depletion strain EC1908 grown in the presence ($FtsN^+$) or absence ($FtsN^–$) of 0.2% L-arabinose. Scale bar: 2µm. **b**. Immunoblot of FtsN using anti-FtsN antiserum from Dr. David S. Weiss[20]. Slight degradation of FtsN during the experiment causes the multiple banding pattern. **c.** Quantification of the blots shows that the total FtsN level after overnight depletion decreases to ~ 45% of the WT cells. Cells were grown in M9-glucose medium overnight at room temperature to $OD_{600}$ = 0.5-0.6.

***Captions for Movies***

**MovieS1:**

Phase contrast time-lapse movie of JXY559 (BW25113 *ftsW*$^{I302C}$) cells growing on 3% agarose gel-pad with M9-glucose minimum medium in the absence of MTSES at room temperature. The movie was recorded every ~3.5 minutes Scale bar: 5 µm.

**MovieS2:**

Phase contrast time-lapse movie of JXY559 (BW25113 *ftsW*$^{I302C}$) cells growing on 3% agarose gel-pad with M9-glucose minimum medium at room temperature. The gel-pad was supplemented with 0.1mM MTSES. The movie was recorded every ~3.5 minutes Scale bar: 5 µm.

**MovieS3-9:**

Epi-fluorescence time-lapse imaging of single FtsW-RFP molecules in M9-glucose, corresponding to Fig 2a, b and Extended Data Figure 6 a-d. Frame rate = 2 frames/second; camera pixel size = 81.25 nm. Scale bar: 0.5 µm.

***Reference:***


1.Neidhardt, F. C., Bloch, P. L. & Smith, D. F. Culture medium for enterobacteria. *J Bacteriol* **119,** 736–747 (1974).

2.Yang, X. *et al.* GTPase activity-coupled treadmilling of the bacterial tubulin FtsZ organizes septal cell wall synthesis. *Science* **355,** 744–747 (2017).

3.Buss, J. *et al.* A multi-layered protein network stabilizes the Escherichia coli FtsZ-ring and modulates constriction dynamics. *PLoS Genet* **11,** e1005128 (2015).

4.Glascock, C. B. & Weickert, M. J. Using chromosomal lacIQ1 to control expression of genes on high-copy-number plasmids in Escherichia coli. *Gene* **223,** 221–231 (1998).

5.Moore, D. A., Whatley, Z. N., Joshi, C. P., Osawa, M. & Erickson, H. P. Probing for Binding Regions of the FtsZ Protein Surface through Site-Directed Insertions: Discovery of Fully Functional FtsZ-Fluorescent Proteins. *J Bacteriol* **199,** 18 (2016).

6.Baba, T. *et al.* Construction of Escherichia coli K-12 in-frame, single-gene knockout mutants: the Keio collection. *Mol Syst Biol* **2,** 2006.0008 (2006).

7.Labun, K., Montague, T. G., Gagnon, J. A., Thyme, S. B. & Valen, E. CHOPCHOP v2: a web tool for the next generation of CRISPR genome engineering. *Nucleic Acids Res* **44,** W272–6 (2016).

8.Hamilton, C. M., Aldea, M., Washburn, B. K., Babitzke, P. & Kushner, S. R. New method for generating deletions and gene replacements in Escherichia coli. *J Bacteriol* **171,** 4617–4622 (1989).

9.Schneider, C. A., Rasband, W. S. & Eliceiri, K. W. NIH Image to ImageJ: 25 years of image analysis. *Nat Methods* **9,** 671–675 (2012).

10.Thévenaz, P., Ruttimann, U. E. & Unser, M. A pyramid approach to subpixel registration based on intensity. *IEEE Trans Image Process* **7,** 27–41 (2008).

11.Ovesný, M., Křížek, P., Borkovec, J., Svindrych, Z. & Hagen, G. M. ThunderSTORM: a comprehensive ImageJ plug-in for PALM and STORM data analysis and super-resolution imaging. *Bioinformatics* **30,** 2389–2390 (2014).

12.Sbalzarini, I. F. & Koumoutsakos, P. Feature point tracking and trajectory analysis for video imaging in cell biology. *J Struct Biol* **151,** 182–195 (2005).

13.Lyu, Z., Coltharp, C., Yang, X. & Xiao, J. Influence of FtsZ GTPase activity and concentration on nanoscale Z-ring structure in vivo revealed by three-dimensional Superresolution imaging. *Biopolymers* **105,** 725–734 (2016).

14.Burman, L. G. & Park, J. T. Molecular model for elongation of the murein sacculus of Escherichia coli. *Proc Natl Acad Sci USA* **81,** 1844–1848 (1984).

15.Egan, A. J. F. & Vollmer, W. The stoichiometric divisome: a hypothesis. *Front Microbiol* **6,** 455 (2015).

16.Cho, H. *et al.* Bacterial cell wall biogenesis is mediated by SEDS and PBP polymerase families functioning semi-autonomously. *Nat Microbiol* **1,** 16172 (2016).

17.Datsenko, K. A. & Wanner, B. L. One-step inactivation of chromosomal genes in Escherichia coli K-12 using PCR products. *Proc Natl Acad Sci USA* **97,** 6640–6645 (2000).

18.Liu, B., Persons, L., Lee, L. & de Boer, P. A. J. Roles for both FtsA and the FtsBLQ subcomplex in FtsN-stimulated cell constriction in Escherichia coli. *Mol Microbiol* **95,** 945–970 (2015).

19.Gerding, M. A. *et al.* Self-enhanced accumulation of FtsN at Division Sites and Roles for Other Proteins with a SPOR domain (DamX, DedD, and RlpA) in Escherichia coli cell constriction. *J Bacteriol* **191,** 7383–7401 (2009).

20.Tarry, M. *et al.* The Escherichia coli cell division protein and model Tat substrate SufI (FtsP) localizes to the septal ring and has a multicopper oxidase-like structure. *J Mol Biol* **386,** 504–519 (2008).

21.Guyer, M. S., Reed, R. R., Steitz, J. A. & Low, K. B. Identification of a sex-factor-affinity site in E. coli as gamma delta. *Cold Spring Harb Symp Quant Biol* **45 Pt 1,** 135–140 (1981).

22.Bernhardt, T. G. & de Boer, P. A. J. The Escherichia coli amidase AmiC is a periplasmic septal ring component exported via the twin-arginine transport pathway. *Mol Microbiol* **48,** 1171–1182 (2003).

23.Guzman, L. M., Belin, D., Carson, M. J. & Beckwith, J. Tight regulation, modulation, and high-level expression by vectors containing the arabinose PBAD promoter. *J Bacteriol* **177,** 4121–4130 (1995).

24.Liang, H. *et al.* Metabolic labelling of the carbohydrate core in bacterial peptidoglycan and its applications. *Nat Commun* **8,** 15015 (2017).

25.Hale, C. A. & de Boer, P. A. J. ZipA is required for recruitment of FtsK, FtsQ, FtsL, and FtsN to the septal ring in Escherichia coli. *J Bacteriol* **184,** 2552–2556 (2002).

26.Mercer, K. L. N. & Weiss, D. S. The Escherichia coli cell division protein FtsW is required to recruit its cognate transpeptidase, FtsI (PBP3), to the division site. *J Bacteriol* **184,** 904–912 (2002).

27.Cho, H., Uehara, T. & Bernhardt, T. G. Beta-lactam antibiotics induce a lethal malfunctioning of the bacterial cell wall synthesis machinery. *Cell* **159,** 1300–1311 (2014).

28.Ducret, A., Quardokus, E. M. & Brun, Y. V. MicrobeJ, a tool for high throughput bacterial cell detection and quantitative analysis. *Nat Microbiol* **1,** 16077 (2016).

29.Sjodt, M. *et al.* Structure of the peptidoglycan polymerase RodA resolved by evolutionary coupling analysis. *Nature* **556,** 118–121 (2018).

30.Kelley, L. A., Mezulis, S., Yates, C. M., Wass, M. N. & Sternberg, M. J. E. The Phyre2 web portal for protein modeling, prediction and analysis. *Nat Protoc* **10,** 845–858 (2015).

31.Rowlett, V. W. & Margolin, W. 3D-SIM super-resolution of FtsZ and its membrane tethers in Escherichia coli cells. *Biophys J* **107,** L17–20 (2014).

32.Kusumi, A., Sako, Y. & Yamamoto, M. Confined lateral diffusion of membrane receptors as studied by single particle tracking (nanovid microscopy). Effects of calcium-induced differentiation in cultured epithelial cells. *Meth Enzymol* **65,** 2021–2040 (1993).
